# A neuroanatomically grounded optimal control model of the compensatory eye movement system

**DOI:** 10.1101/617365

**Authors:** P.J. Holland, T.M. Sibindi, M. Ginzburg, S. Das, K. Arkesteijn, M.A. Frens, O. Donchin

## Abstract

We present a working model of the compensatory eye movement system. We challenge the model with a data set of eye movements in mice (n=34) recorded in 4 different sinusoidal stimulus conditions with 36 different combinations of frequency (0.1-3.2 Hz) and amplitude (0.5-8°) in each condition. The conditions included vestibular stimulation in the dark (vestibular-ocular reflex, VOR), optokinetic stimulation (optokinetic reflex, OKR), and two combined visual/vestibular conditions (the visual-vestibular ocular reflex, vVOR, and visual suppression of the VOR, sVOR). The model successfully reproduced the eye movements in all conditions, except for minor failures to predict phase when gain was very low. Most importantly, it could explain the non-linear summation of VOR and OKR when the two reflexes are activated simultaneously during vVOR stimulation. In addition to our own data, we also reproduced the behavior of the compensatory eye movement system found in the existing literature. These include its response to sum-of-sines stimuli, its response after lesions of the nucleus prepositus hypoglossi or the flocculus, characteristics of VOR adaptation, and characteristics of drift in the dark. Our model is based on ideas of state prediction and forward modeling that have been widely used in the study of motor control. However, it represents one of the first quantitative efforts to simulate the full range of behaviors of a specific system. The model has two separate processing loops, one for vestibular stimulation and one for visual stimulation. Importantly, state prediction in the visual processing loop depends on a forward model of residual retinal slip after vestibular processing. In addition, we hypothesize that adaptation in the system is primarily adaptation of this model. In other words, VOR adaptation happens primarily in the OKR loop.

## Introduction

Optimal control is a widely used paradigm in current models of motor behavior (Frens and Donchin, 2009; Haar and Donchin, 2019 [preprint]; Parrell et al., 2019 [preprint]; Shadmehr and Krakauer, 2008). Optimal control suggests that the motor system operates in a “full feedback” mode: generating motor commands in response to the best guess regarding the current situation as opposed to using a pre-defined plan (Todorov and Jordan, 2002). However, it has proved very difficult to build optimal control models that make specific predictions for real, physiological motor circuits. In this paper, we address this gap by building a working quantitative model (Fig 1) of the compensatory eye movement system (CEM) starting from the ideas developed in the Frens and Donchin state predicting feedback control (SPFC) scheme (Frens and Donchin, 2009).

**Figure 1.**
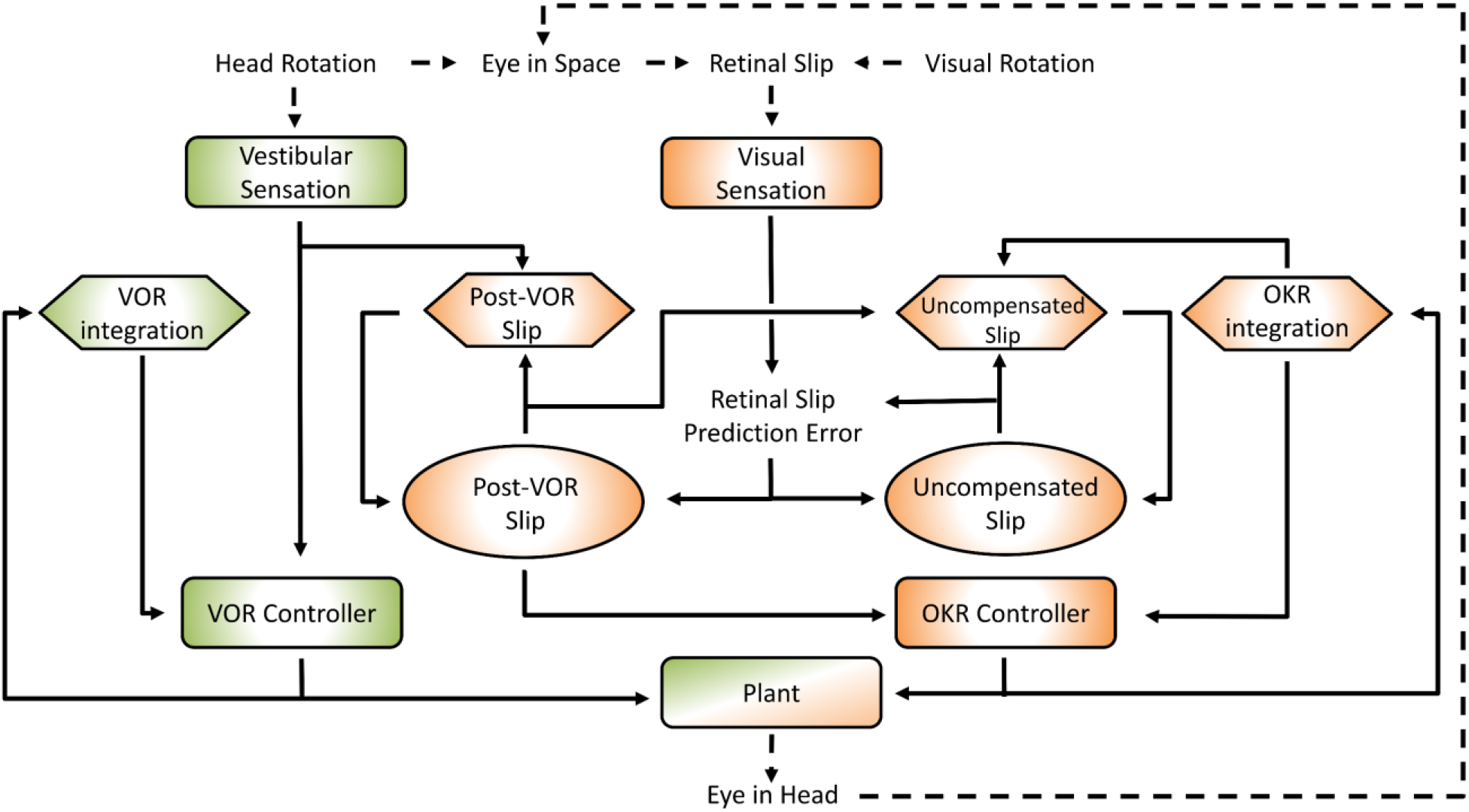
General layout of the model. Green areas are vestibular, orange areas are optokinetic. Hexagons represent Forward Models, ellipses are State Estimators. Dashed arrows indicate processes in the real world, solid arrows are neural processes. Details of the model are specified in the text and supplementary material.

Compensatory eye movement is a general term for several reflexes whose goal is to maintain a stable image on the retina during movements of the head by moving the eyes in the opposite direction (Delgado-García, 2000). In other words, these reflexes serve to reduce retinal slip (movement of the visual image across the retina). The CEM system has a number of properties that make it a popular candidate for quantitative modeling of sensorimotor processes (Lisberger, 2009). First, its goal, minimizing retinal slip, is clear and invariant over time. Second, the dynamics of the system as a whole are close to linear. Third, the output only has three degrees of freedom. Moreover, horizontal CEM can be isolated from the other two degrees of freedom and treated as a system with a single degree of freedom. This is commonly done in the experimental literature, and it is our approach as well. However, since rotations are non-commutative, expanding the model to three dimensions is not trivial.

We chose to model and perform experiments in mice because mice, being afoveate, lack a confounding smooth pursuit system. In afoveate animals like mice, the CEM comprises two reflexes: the vestibulo-ocular reflex (VOR) uses vestibular input to predictively compensate retinal slip and the optokinetic reflex (OKR) is driven by the retinal slip itself. The two reflexes have roughly complementary properties: the OKR performs well in low velocities and the VOR works well at high frequencies. The existence of these reflexes allows accurate compensation of the retinal slip velocity in normal behavior. However, a challenge for any model of the CEM is to explain the non-linear interaction between VOR and OKR. In many conditions, the combined action, with a gain of almost exactly one, is much less than the sum of the two reflexes driven separately.

One possible explanation for their non-linear summation is that the OKR is capable of predicting the retinal slip that remains after the VOR. This is at the core of the model that we present. The current model is essentially hierarchical, with the vestibular and the visual components of the CEM handled in two distinct loops (see Fig 1). This is close to the traditional view of CEM which also incorporates two, more or less separate, mechanisms for the VOR and OKR (Wakita et al., 2017). The VOR operates in a partially open-loop fashion with feedback used to drive only the forward model of the eye without modifying processing of the vestibular state itself. The OKR loop, on the other hand, incorporates forward models of the eye, the visual input, and also the VOR system. That is, the OKR not only predicts current retinal slip based on models of the environment and the eye movements, it also incorporates a model of the residual retinal slip that remains after the actions of the VOR loop.

We wanted a model that reproduces the main characteristics of mouse vVOR (rotation of the animal in the light, providing simultaneous visual and vestibular stimulation), without needing to carefully tweak the model parameters. Furthermore, the same set of parameters should then result in good predictions of responses in VOR, OKR and additional conditions, i.e. suppressed VOR (sVOR; simultaneous rotation of the animal and its visual surroundings), and responses to sum-of-sines (SOS) stimuli. Furthermore, in order to test the relation between the different pieces of the model and the underlying anatomy, lesions in specific parts of the model should mimic actual lesions in the associated brain structures.

We also modeled adaptation. We postulate that the primary adaptation of the CEM system is in the OKR part of the system. This is consistent with experimental findings (as reviewed in the discussion) and also with our hypothesis that the OKR loop is more dependent on forward model prediction than the VOR Thus, adaptation of the CEM system (at least to first approximation) is mostly adaptation of the OKR model of VOR inaccuracies (Fig 1; Post-VOR Slip). An additional test of our model is that it should be possible to set the value of ζ adaptively, thus mimicking VOR adaptation

Finally, in order to compare our model to data, we collected from mice in a large set of conditions (VOR, OKR, vVOR, sVOR, SOS), frequencies and amplitudes. Such a data set was lacking in the literature so that our contribution in this work, beyond a model that fits all existing data, is a comprehensive data set showing OKR behavior across a complete array of stimuli.

## Materials and Methods

### Model

The model was implemented in Matlab (version 2016a; The MathWorks, Natick, MA, USA) and calculations were performed via matrix multiplication with a time step of 1 ms. Details are provided in the Supplementary Material.

**VOR:** The mouse VOR uses vestibular input from the semi-circular canals (labyrinth) to compensate head movement (Delgado-García, 2000). Vestibular afferents from the labyrinth project directly to VN with a small delay (2ms; Sohmer et al., 1999). Their activity accurately reflects head velocity at high frequencies but not at low frequencies (Robinson, 1981) due to filtering properties of the vestibular labyrinth (Yang and Hullar, 2007). Thus, in modeling VOR, the processing is quite simple (green areas in Fig 1). Since the system has no access to the actual head velocity, we use the vestibular signal as an approximation of the head velocity. Neither system dynamics nor the oculomotor command affect head dynamics. Note, therefore, that this model currently does not distinguish between active and passive head movements, i.e. it does not incorporate efference copy or proprioceptive information about head movement.

The job of the second part of the control loop is to estimate the retinal slip that will be uncompensated by the VOR (Post-VOR Slip) and then compensate for it. Post-VOR slip arises from two sources: from changes in the velocity of the visual stimulus and from head movements not compensated by the VOR. These signals represent the predicted retinal slip for which the OKR needs to correct. The combination of this predicted retinal slip combined with an estimate of how much the OKR is moving the eye, gives the OKR’s forward model prediction of uncompensated retinal slip (right orange hexagon in Fig 1).

**OKR:** In the mouse, the OKR originates in velocity sensitive neurons of the retina, which project through the Accessory Optic System (AOS) and Nucleus Reticularis Tegmenti Pontis (NRTP) to the vestibular nucleus (VN) and the vestibulo-cerebellum (Gerrits et al., 1984; Glickstein et al., 1994; Langer et al., 1985). The VN output is sent to the brainstem nuclei, which drive the extra-ocular muscles. In the case of horizontal eye movements, these are the abducens nucleus (Ab), the oculomotor nucleus (OMN) and nucleus prepositus hypoglossi (NPH; Büttner-Ennever and Büttner, 1992).The OKR has a species-dependent response delay of 70-120 ms (van Alphen et al., 2001; Collewijn, 1969; Winkelman and Frens, 2006) primarily caused by the visual processing in the pathway from retina to VN (Graf et al., 1988). The retinal afferents saturate at high velocities (Oyster et al., 1972; Soodak and Simpson, 1988), causing non-linearities in the OKR in this range (van Alphen et al., 2001; Collewijn, 1969). Thus, the OKR is ineffective in compensating high velocity (and thus often high frequency) visual stimuli.

In our model, the OKR system models the effect of VOR as a linear correction for the sensed head velocity. That is, it assumes that VOR compensates for some fraction of the head movement. Thus, our forward model estimate of movement of the visual surrounding (Post-VOR Slip; left orange hexagon in Fig 1) will be updated by a factor proportional to head acceleration (See also Eq 29 in Supplementary text). The specific constant of proportionality, *ζ*, is discussed in the section on VOR adaptation below.

As one can see in Figure 1, state estimation produces estimates of both Post-VOR slip, and uncompensated retinal slip (oval boxes). Post-VOR slip is retinal slip after VOR compensation and uncompensated slip is that remaining after the action of both systems. The state estimator approximates a Kalman filter, where gain was chosen by hand to match the data instead of being set at the Kalman gain (see Supplementary Material, Eq 42). Thus, through the model architecture, vestibular input only affects our estimate of the head velocity, and retinal input affects both our estimate of retinal slip and our estimate of uncompensated retinal slip.

#### VOR adaptation

VOR adaptation occurs when gaze consistently fails to compensate head movement (Blazquez et al., 2004; Schonewille et al., 2010; Shin et al., 2014). In a laboratory environment, a rotating visual environment can lead such failure (as described in the Methods below). This causes persistent changes in the VOR, such that retinal slip is reduced in the new situation. In our model, such a mismatch would affect the proportionality constant ζ. This is because the OKR system’s assumption that retinal slip is the result of inaccuracies in the VOR loop (see Supplementary Material).

#### Parameters

In the model only a few parameters were set to match the data. They were set to match data in the vVOR condition and then the same parameters were used for all conditions. Most variables were either taken from literature, or experimentally derived by us in separate experiments. Interestingly, it turned out that the model produced very similar behavior across a wide range of values for most parameters although it was sensitive to a few parameters (see table 1). As much as possible, parameters were determined from the literature or from our own data. For example, we determined the maximum VOR and OKR gains from our own data. We used the response to high frequency stimulation to set the maximum gain of the VOR in the model and the response to low velocity stimulation to set the maximum gain of the OKR in the model. The form of the non-linearity in retinal slip was fit to published results (Oyster et al., 1972; Soodak and Simpson, 1988). On the other hand, the filter of the vestibular afferents was shaped to achieve the best fit to the data. Ultimately, the filter that fit our data best was also compatible with the literature. We used a first order high pass filter with a time constant of 4 s (Yang and Hullar, 2007). Similarly, drift velocity and VOR adaptation speed were fit to data and later found to be compatible with the literature (Schonewille et al., 2010; Stahl et al., 2006).

**Table 1.** Overview of all parameters used in the model, their values, the equations they are used (described in Supplementary Material), and a short description of their meaning. The last two columns describe whether they are critical, and how they were set. We determined how critical the parameters were, by varying them over an order of magnitude, and observing the changes in results.

|  | Value | Eq. | Meaning | Is it critical? | How was it set? |
| --- | --- | --- | --- | --- | --- |
| $dt$ | 1 ms | | Time step | | |
| $T_p$ | 0.5 sec | 1, 2,<br>15,<br>17,<br>24,<br>33,<br>38 | Leaky integrator time constant for motor nuclei | No | (Stahl and Simpson, 1995; Stahl et al., 2015) |
| $T_v$ | 4 sec | 3, 15 | Low pass filter constant for the vestibular inputs | Yes | Fit to data. Close to value found for actual vestibular afferents by Yang and Hullar 2007 (3 sec). |
| $\delta_v$ | 2 ms | 5, 20 | Vestibular sensory delay | No | |
| $a_v$ | 0.1 | 6, 16 | Vestibular sensory noise proportionality constant | No | Middle of the stable range |
| $R_{\max}$ | 0.65 deg/sec | 8 | Retinal saturation | Yes | Oyster et al, 1972 |
| $\delta_R$ | 70 ms | 9,<br>20,<br>31 | Visual processing delay | Yes | Van Alphen, 2001 |
| $a_R$ | 0.1 | 10,<br>16 | Visual sensory noise proportionality constant | No | Middle of the stable range. |
| $a_u$ | 0.1 | 15,<br>16 | Motor noise | No | Middle of the stable range |
| $Z$ | -0.6 | 29,<br>33,<br>38,<br>48 | Assumed VOR inaccuracy | No | Fit to match VOR performance in the dark. |
| $\kappa_V$ | 1 | 22,<br>42 | Kalman gain of vestibular input | No | Set so VOR is not eliminated by floccular lesion |
| $\kappa_T$ | 0.05 | 30,<br>42 | Kalman gain for the effect of retinal slip prediction error on assumed external motion (post-VOR retinal slip) | No | Fit to data |
| $\kappa_{R,k}$ | 0.05 | 31,<br>35,<br>42 | Kalman gain for the effect of retinal slip prediction error on estimate of uncompensated retinal slip | No | Fit to data |
| $\gamma$ | $e^{\frac{1}{150}}$ | 44 | Discount parameter for cost function | No | Fit to produce credible drift in the dark |
| $\theta$ | 2 | 44 | Weight of position factor in cost function | No | Fit to produce credible drift in the dark |
|  | 100,000 |  | Number of terms kept in infinite cost function sum | No | Arbitrary |

### Animals

In order to test the model we recorded CEM in 13 C57Bl/6J mice (Charles River, Wilmington, MA, USA). We employed four different paradigms i.e. OKR, VOR, sVOR, and vVOR and in each condition we tested a wide range of frequency and amplitude combinations. Details on the experiments are described in the Supplementary Material. Additionally, we measured the drift of the eye back to a central position in the dark (N=6) and the rate of adaptation of the VOR (N=7), full details of the methods are described in the supplementary material.

Prior to all eye movement recordings, mice underwent surgery to prepare them for head fixation and were allowed sufficient time to recover, details are provided in the supplementary material and the full procedure is described in van Alphen (2009).

During an experimental session, mice were immobilized by placing them in a plastic tube with the head protruding and the head fixation attached to the turntable with the eye in the central position. Eye movements were recorded via an infra-red video system (Iscan ETL-200, Iscan, Burlington, MA, USA) at a frequency of 120 Hz. Visual stimuli were presented using a modified projector (Christie Digital Systems, Cypress, CA, USA) displaying a panoramic field of 1592 green dots on virtual sphere fully surrounding the animal. Rotation of the sphere around the vertical axis provided the moving stimuli. Vestibular stimulation was provided via a motorized turntable Mavilor-DC motor 80 (Mavilor Motors S.A., Barcelona, Spain) on which the mouse and eye movement recording system were mounted. Further details are provided in the supplementary material and a schematic representation of the stimulus and eye movement recording apparatus in Figure 2.

**Figure 2.**
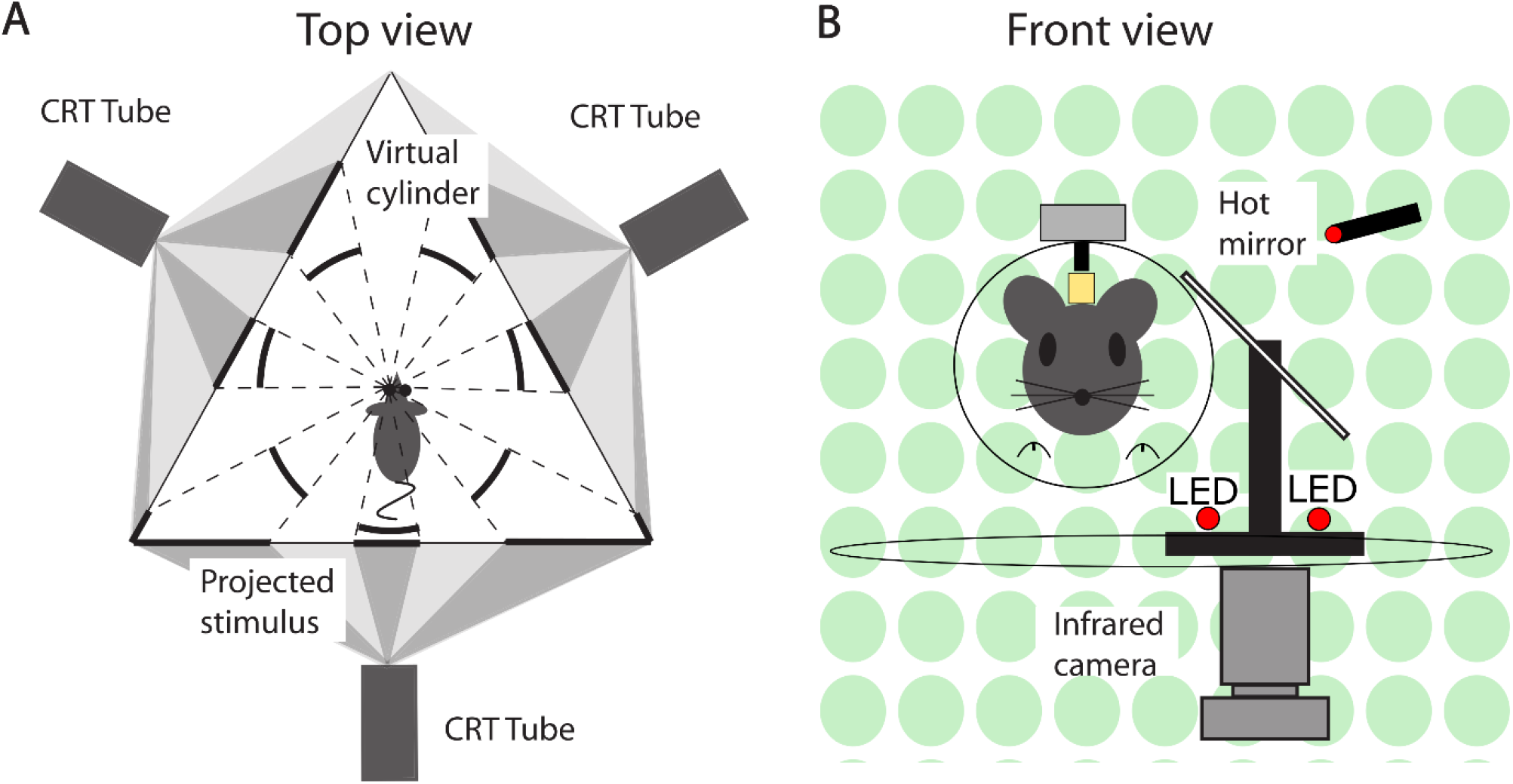
Schematic representation of the experimental setup. (A) Top view. A mouse in the setup, with its left eye in the center and surrounded by three screens on which the visual stimuli are projected. The visual stimuli were programmed and displayed in such a way that from the point of view of mouse it appeared as a virtual sphere. (B) Front view. A mouse placed in front of a hot mirror, which enabled the infrared camera underneath the table to record the eye movements.

The VOR adaptation experimental paradigm consisted of an identical stimulus setup with the animal undergoing 6 VOR trials (1 min duration, 1 Hz, 5°) to measure the gain alternating with 5 sVOR trials (5 min duration, 1 Hz, 5°) to induce adaptation.

### Data Analysis

Every mouse was tested once in each condition, and each stimulus consisted of at least 5 cycles. Full details of the analysis details are provided in the supplementary material. Briefly, following filtering and removal of fast phase eye movements gain and phase data was calculated by a Bayesian fitting procedure in OpenBugs (Version 3.2.3, http://www.openbugs.net, [Lunn et al., 2009]) and Matlab curve fitting routines, for single sinusoid stimuli and for SoS stimuli respectively. The Matlab code and data required for replication of the analysis presented in this paper is available on the Open Science Framework website (https://osf.io/feq7c/).

## Results

### Responses to sinusoidal stimulation

The behavioral data that we present are in agreement with the values that have been previously published for the C57BL/6 mouse strain (van Alphen et al., 2010; Faulstich et al., 2004; Schonewille et al., 2011; Stahl et al., 2000). The VOR (Fig 3) in the dark responded to high frequency stimulation, and the OKR (Fig 4) was mainly active in response to low velocity stimuli (van Alphen et al., 2001). The vVOR (Fig 5) was more or less veridical over the whole stimulus range while suppression in the sVOR (Fig 6) paradigm mainly happened at low frequency/velocity conditions.

**Figure 3.**
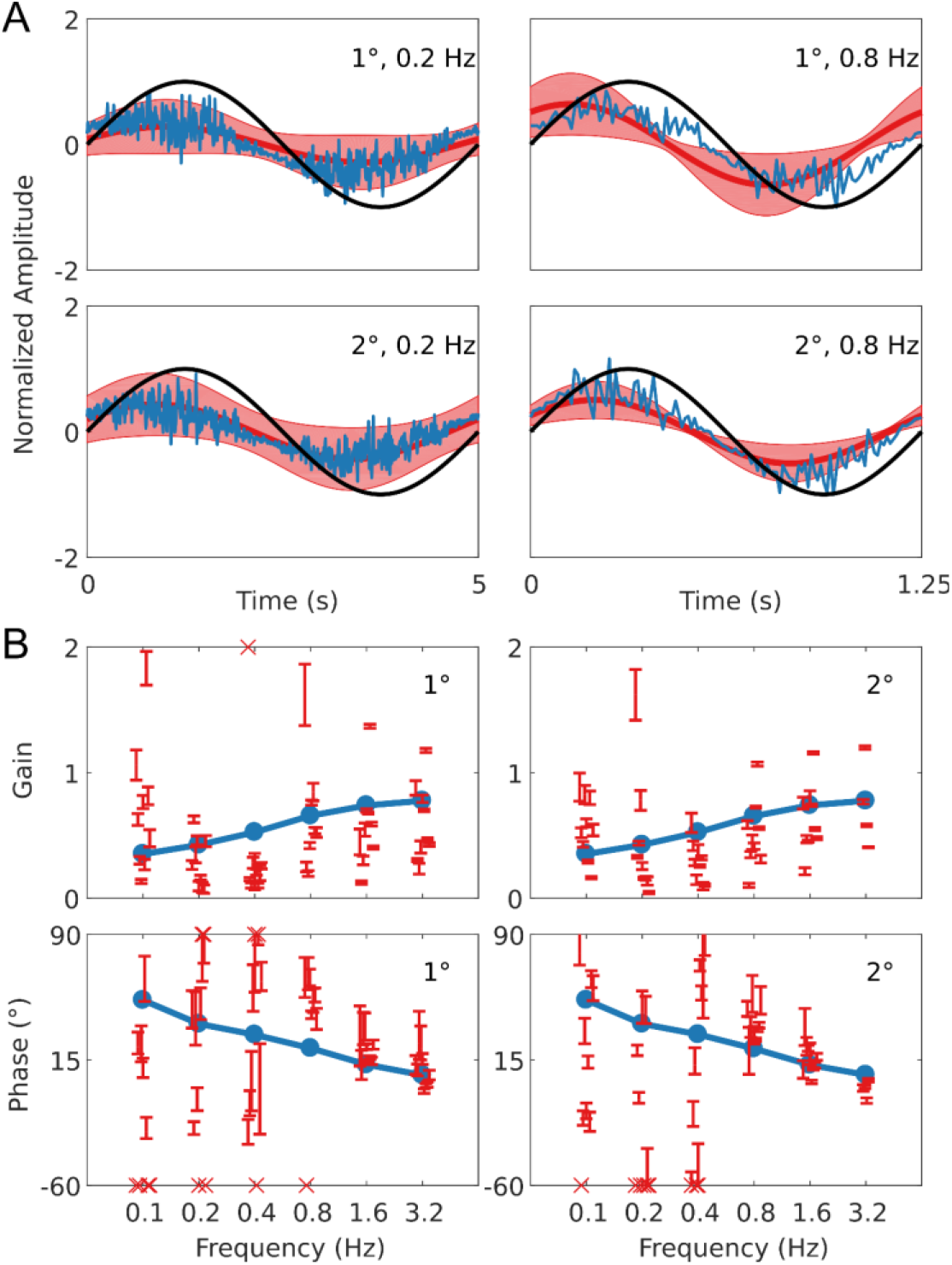
Summary of VOR data and simulation. In panel A the upper row displays results for 1° stimuli, the lower row for 2° stimuli. The panels show the stimulus in black (left: 0.2Hz; right 0.8Hz), with the simulated response (blue) and the mean measured responses (red). Shaded red regions represent the standard deviation (SD) of the population. Panel B are Bode plots for Gain (top panels) and Phase (bottom panels) for the simulated response (blue), individual mice with SD (red dots).Crosses in the Phase plots indicate conditions where the response was too small to determine a phase reliably. The left and right sides of Panel B represent bode plots for 1° stimuli and 2° stimuli respectively. Other stimulus conditions fitted equally well.

**Figure 4.**
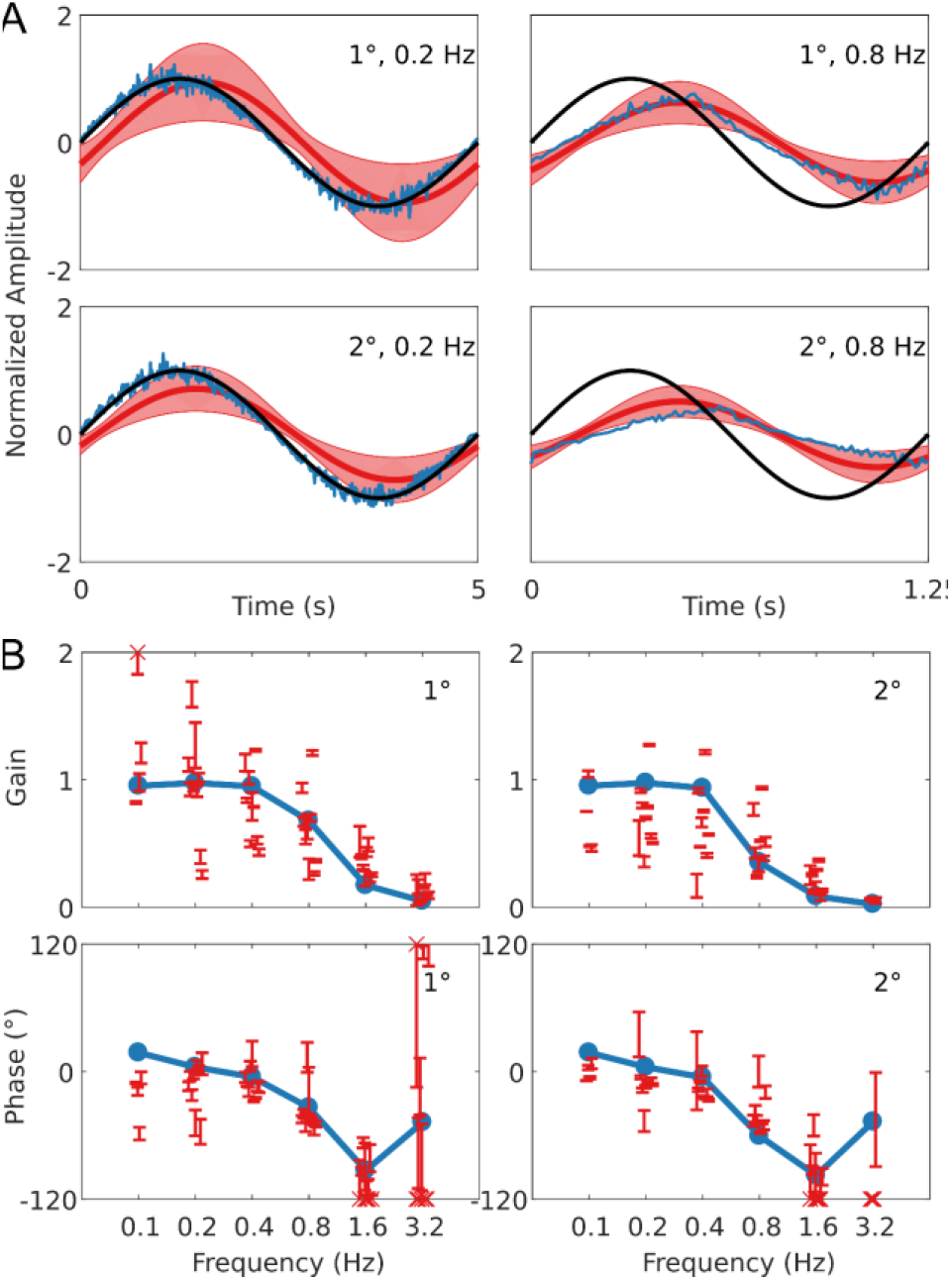
Summary of OKR data and simulation. This figure follows the format of Fig 3. Note that the phase response of stimuli with Gains < 0.25 could often not reliably be determined.

**Figure 5.**
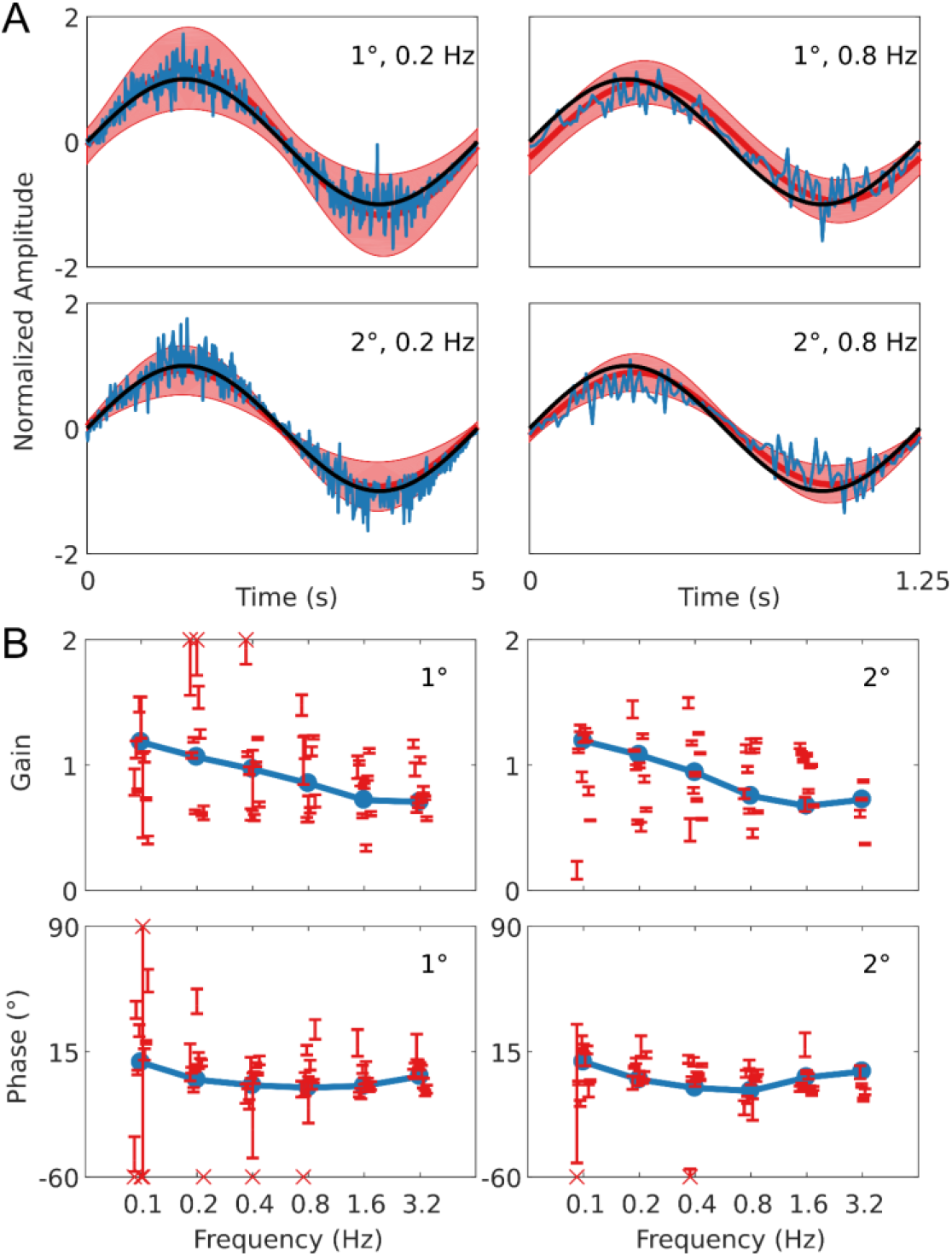
Summary of vVOR data and simulation. This figure follows the format of Fig 3. Across the whole frequency range tested in both amplitudes there was a very good match of model to experimental data.

**Figure 6.**
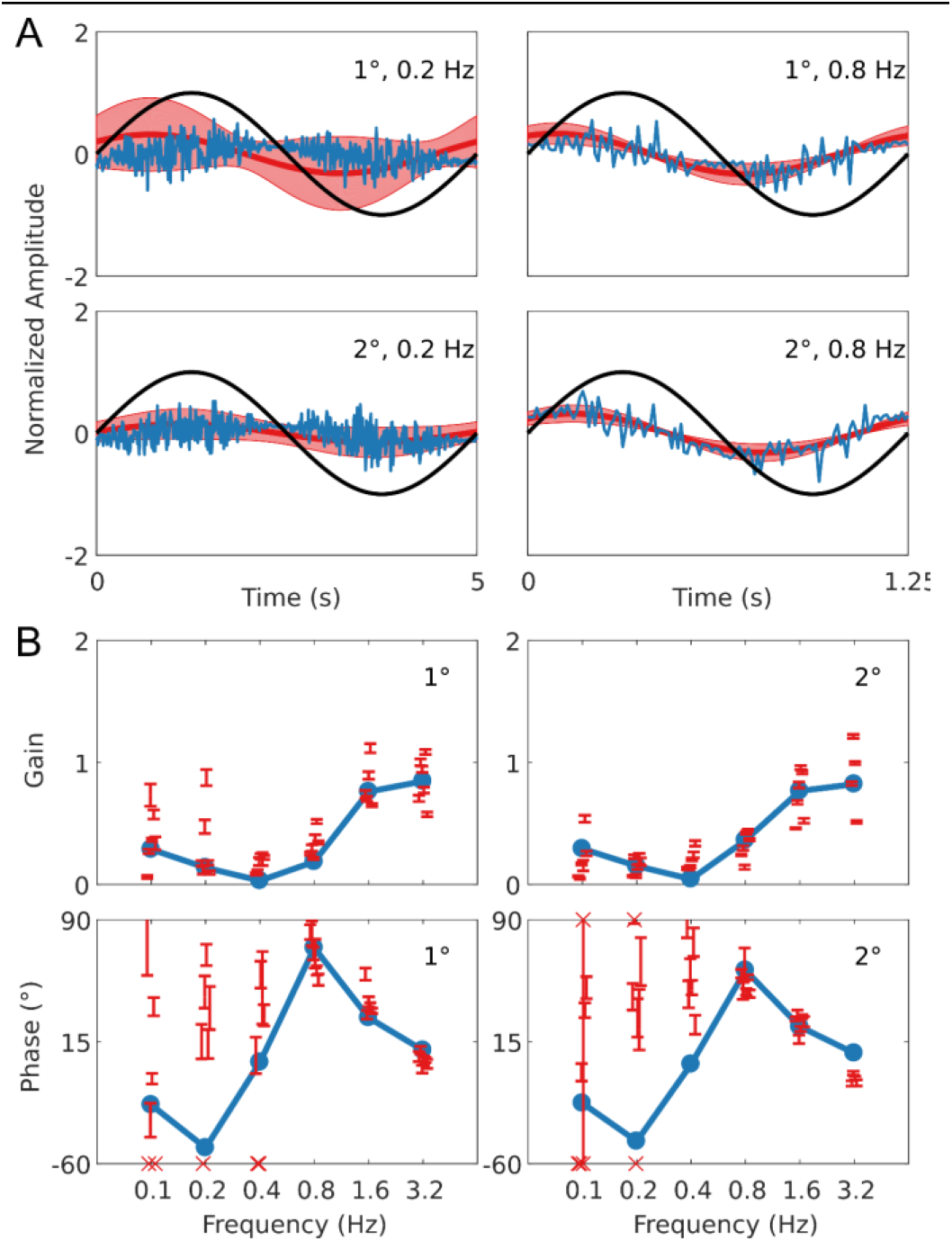
Summary of sVOR data and simulation. This figure follows the format of Fig 3. Note that hase response of stimuli with Gains < 0.25 could often not reliably be determined. The pattern of the response in the behavioral data is clearly captured by the simulation.

In Figure 3 we show a comparison of experimental and simulated VOR. We see that there is a good match between simulation and average experimental response over the whole stimulus range. First, a high gain at high frequencies and lower gain at low frequencies is clearly observable. Furthermore, we see a phase lead at low frequencies which diminishes with increasing stimulus frequency.

Figure 4 follows the same format as Figure 3 but compares simulation to experimental results for the OKR response. The simulation nicely predicts the main features of the OKR response. The gain decreases and the phase lag increases with increasing stimulus velocity.

Figure 5 shows how well simulations predict experimental data for combined visual and vestibular stimulation (vVOR). In both the simulation and experimental data, we observe high gain and almost no phase lead or lag between response and stimulus. These results show that VOR and OKR have complementary results, which allows the combined system to produce excellent compensation of the retinal slip.

Figure 6 depicts how the model fits experimental data generated during sVOR – suppression of the VOR response with visual input. The response in high frequencies looks very similar to that in VOR because OKR is not responsive in high frequencies (see Fig. 4), and hence cannot suppress vestibular triggered response. At low frequencies, there is a very small response, because VOR has low gain and is further suppressed by OKR. At these low frequencies, where the gain is low and variable, the model systematically misrepresents the phase of the eye movement.

In order to examine the overall quality of fit in each of the four experimental conditions above, we calculated the Z-scores of the overall fitting quality. These are displayed in Figure 7. Note how the overall fit quality is good (“cool” colors in the heat map), with some poorer fits in the low frequency/high amplitude range of the sVOR condition. Because of the low amplitudes and high variability, the phase offset of the model at the lower frequencies does not lead to large Z-scores.

**Figure 7.**
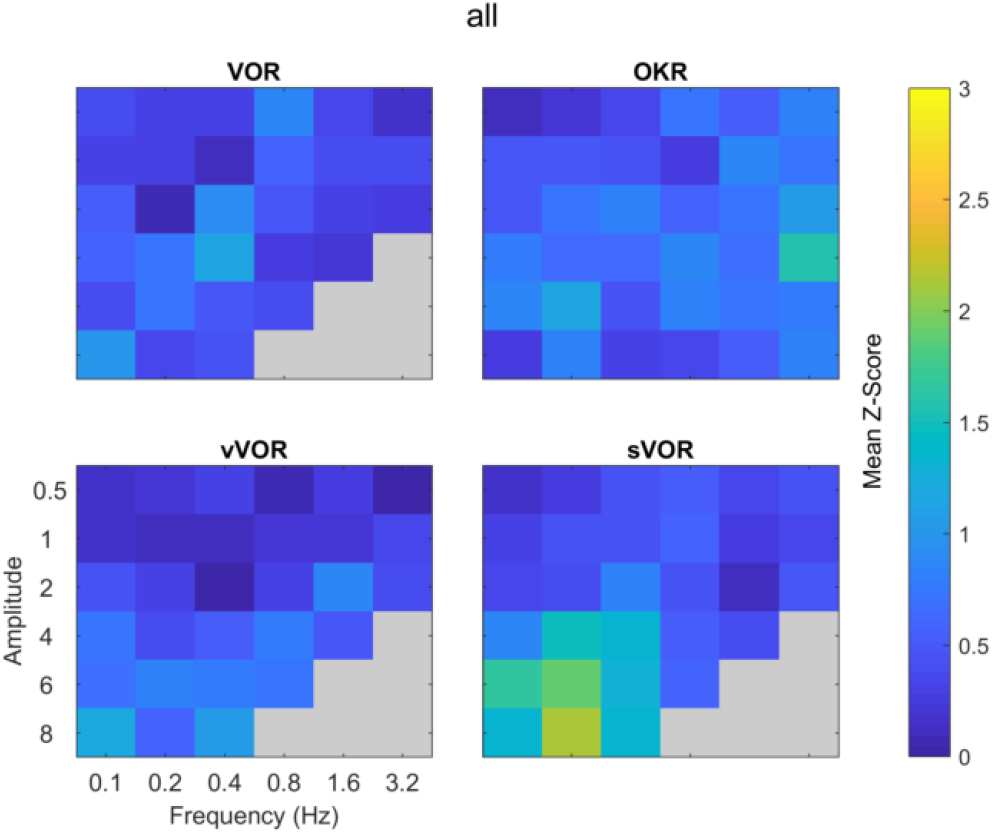
Summary of comparison of model and data for all amplitude and frequency combinations. The four panels depict the degree to which the model response matched the experimental data for the four conditions. The degree of similarity is expressed in terms of the number of standard deviations the model response was away from the mean behavioral response, cooler colors indicate a closer match. Across all amplitude and frequencies tested the model reproduces the experimental data well, with the possible exception of high amplitude, low frequency sVOR. Grey regions indicate conditions not measured in the experimental data

In addition to the comparison of model and data in terms of Z-scores (Fig 7), we also used the Bayesian estimates to generate probabilities for the model response falling outside the range of the behavior of a ‘typical mouse’. These tests were carried out for every frequency and amplitude combination and assessed the similarity of gain and phase separately, a combined probability was then generated from the product of these. The results of these tests are presented in the supplementary material. Overall the model responds within the range of a typical mouse for both gain and phase individually and when combined.

### Model Dynamics

The interaction of the different parts of the model in one of the conditions (vVOR, amplitude 2, frequency 0.2 Hz) are shown in Figure 8. The figure shows one cycle of the activity in each of the different areas being modeled during the steady state response to this stimulus. The top two boxes show that in this condition, the head is being rotated but the eyes are moving to keep the retinal slip at 0. The head rotation passes through the system in a feedforward manner to drive the vestibular controller. Additionally, this controller is modulated by knowledge of the eye position and velocity, driven by the forward model integration of the vestibular command. The figure also shows how the head rotation drives an estimate of the retinal slip that would remain uncompensated by the VOR controller. This is labeled post-VOR slip. Post-VOR slip in turn drives the activity of the OKR controller. Note that in this condition, the system estimates that the VOR will over-compensate for the head rotation and the OKR controller generator actually generates a command that is roughly in counter-phase with that of the VOR controller. The success of the vVOR in generating eye movements that fully compensate for the head movement are the result of a balance between the VOR signal and the OKR signal. Without the balancing OKR signal, the gain of the VOR would need to be lowered to achieve veridical tracking, which would compromise the quality of the VOR.

**Figure 8.**
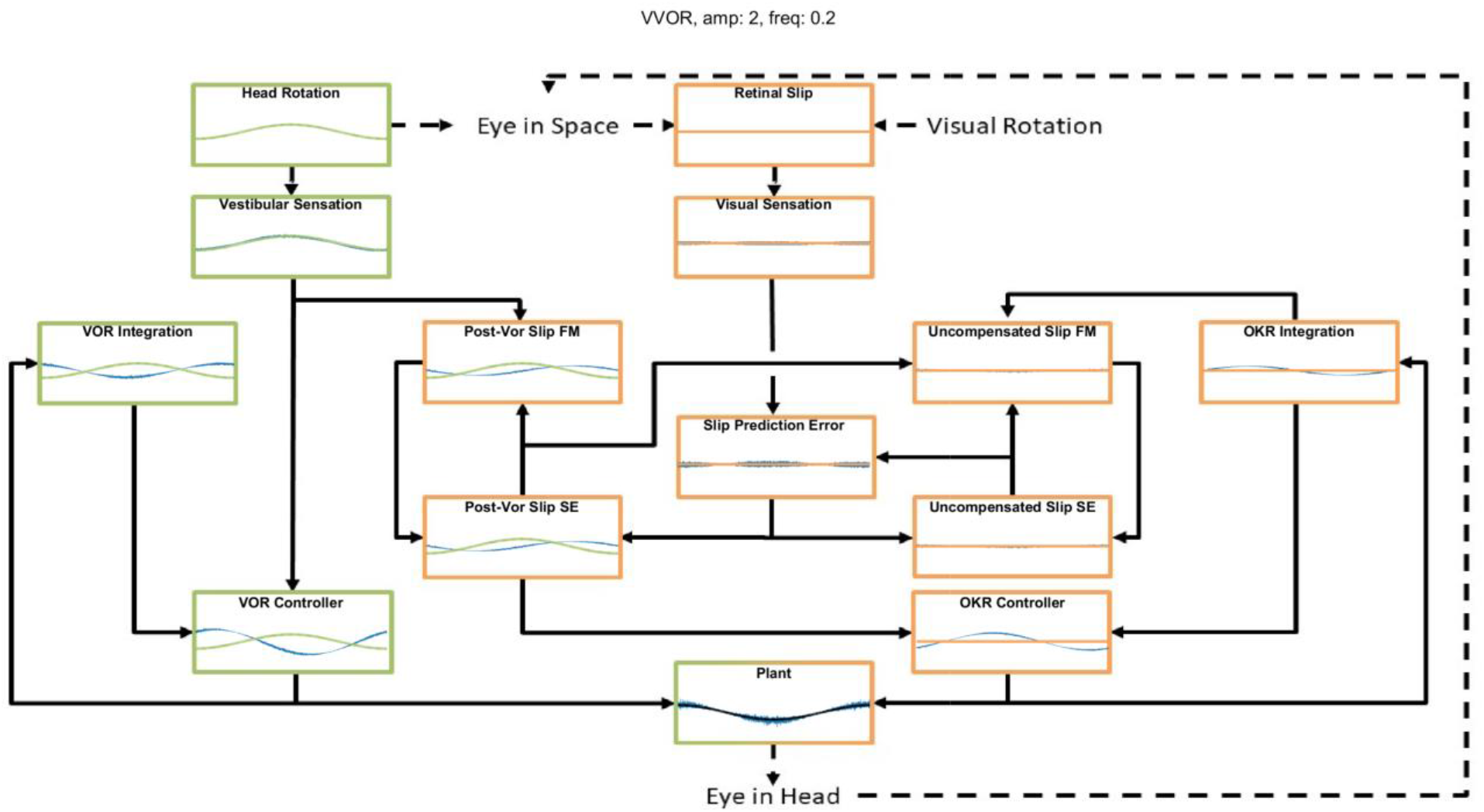
An example of the model dynamics for one cycle of the simulation in the vVOR condition (Stimulation amplitude of 2 degrees at a frequency of 0.2 Hz) at a time by which the system has reached a steady state. The layout matches the model schematic presented in Figure 1. In each box the blue line represents the output of the computation performed, the green or orange line represents the appropriate stimulus, vestibular and visual respectively. In the Supplement we display the full model dynamics for the VOR and OKR conditions in isolation, respectively, for the same frequency and amplitude of stimulation.

Figure 8 also shows that in this situation the OKR system has stabilized, such that retinal slip prediction error is 0. If there were prediction error, generated by either a transient visual or vestibular perturbation, this would drive an increase in the post-VOR slip which would then cause a transient increase in the OKR command to correct for the extra slip. The OKR system thus serves in two complementary roles: it generates a feedforward correction for the inaccuracies of the VOR system (the size of which is learned through adaptation, as described below) and it generates an error driven correction for unexpected retinal slip. The figure thus demonstrates the balance between the VOR command, post-VOR slip and OKR command that are necessary to achieve veridical tracking in the vVOR condition. Figures in the supplementary results show dynamic plots for other stimulus conditions and other frequencies and amplitudes, but it is this interaction which is the key innovation of our model.

### Sum of Sines

When the mouse OKR responds to sum-of-sines (SoS) stimuli, we have previously reported relative gain suppression of the lower of two frequencies in the stimulus. Conversely, in sVOR, results showed gain enhancement in the lower frequency component. In both sVOR and VOR, an overall decrease in phase lead was observed. For more details see (Sibindi et al., 2016). When applying these stimuli to the model, the main pattern of effects is reproduced. Thus, we find qualitatively similar changes in both the relative gain and delay of the constituting frequencies (Fig 9A). Importantly, removal of retinal saturation eliminates the non-linearities expressed in the gain of the response (Fig 9B).

**Figure 9.**
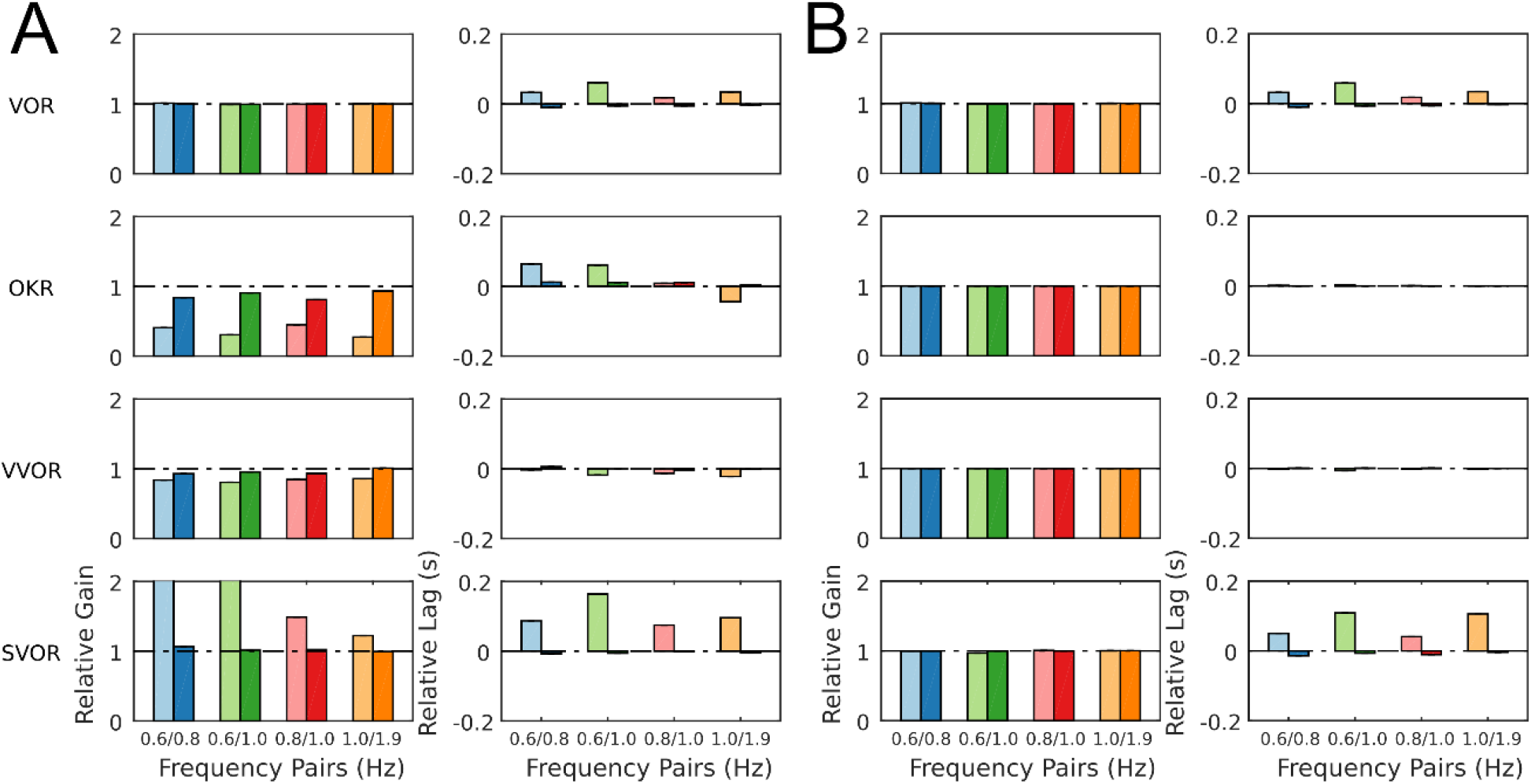
Summary of the model response to Sum of Sines stimulation for the model with normal retinal saturation (A) and with the saturation of retinal input removed (B). The response is described in terms of gains and lags relative to the gain and lag recorded in response to the single frequency component presented in isolation. A linear system will produce only relative gains of 1 and relative delays of 0, indicated by dashed horizontal lines on each plot. The pattern of nonlinearities produced by the full model (A) matches closely the nonlinearities found in behavioral data in response to the same stimuli (Sibindi et al 2016). Figure 6 of Sibindi et al (2016) is reproduced with consent as a supplement to this figure (Figure 7-figure supplement 1). The removal of retinal saturation eliminates the non-linearities expressed in the relative gains of the OKR and sVOR but those expressed in the relative lags of VOR and sVOR remain intact. Please note that the values for relative gain for the 0.6 Hz component of the 0.6/0.8 and 0.6/1.0 Hz Sum of Sines in sVOR (panel A) are greater than 2 and the values are indicated on the bars.

### VOR Adaptation

Perhaps counterintuitively, VOR adaptation occurs as a result of changes in the OKR’s model of VOR. Adaptation modifies the OKR’s prediction of post-VOR slip. Thus, adaptation in our model involved allowing the parameter ζ to vary in response to retinal slip prediction error using gradient descent. As derived in the supplementary material, the gradient is in the direction that decorrelates head acceleration and retinal slip prediction error. The minimum error had a broad basin of attraction. Thus, regardless of the starting value of ζ, it always converged to the same value of −0.6, if the stimulation frequency was kept constant at 1 Hz. The value to which ζ converged depended on stimulus frequency but not amplitude. Nevertheless, for a broad range of frequencies ζ assumed a value around −0.6. The adaptation protocol reduced the gain of the VOR in mice to around 50% of its original value (Fig 10), comparable to that which has been previously described in literature (Schonewille et al., 2011).

**Figure 10.**
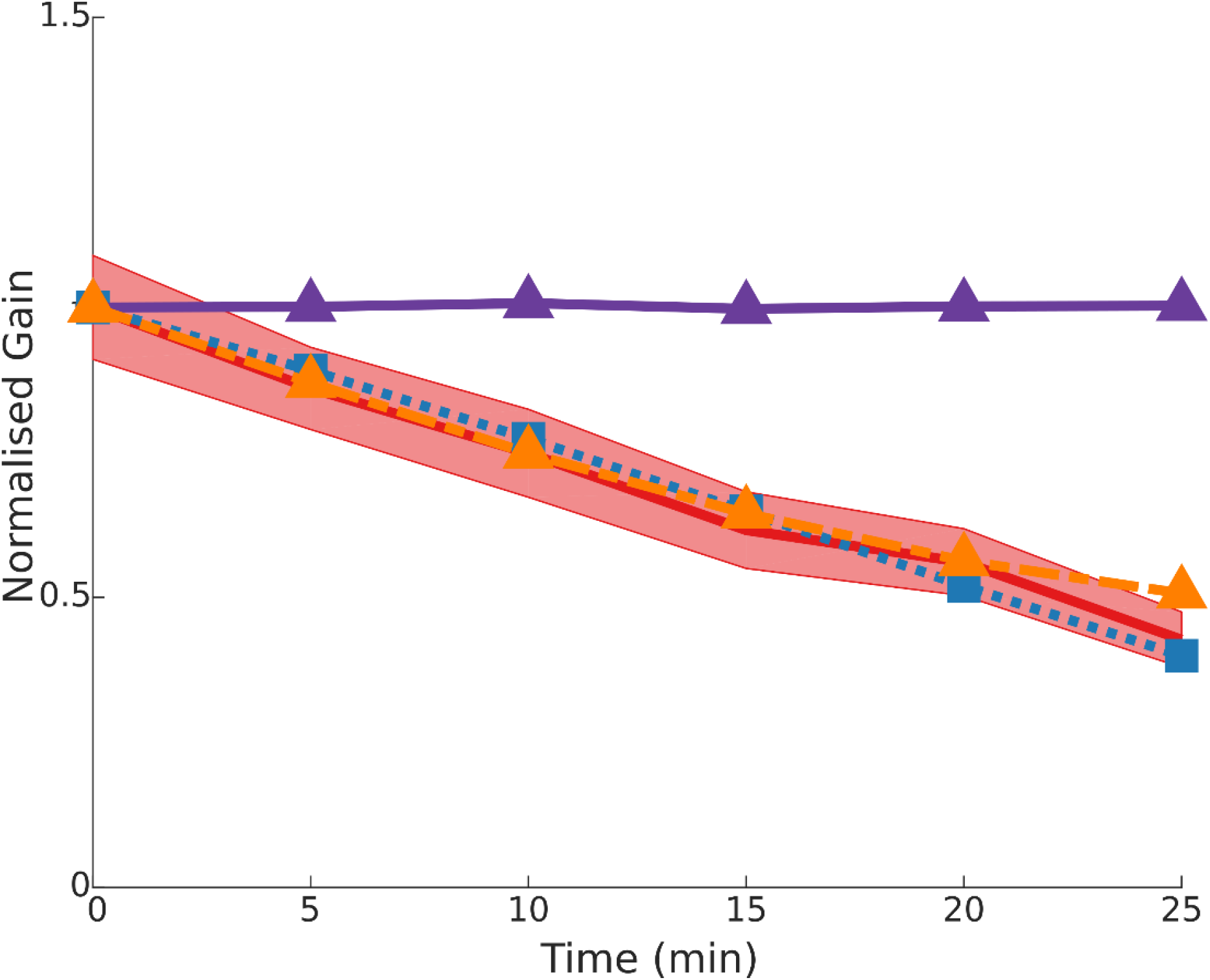
Time course of gain decrease adaptation of the VOR in response to repeated sVOR stimulation. The decrease in gain measured experimentally (red) with confidence limits representing SEM (shaded region) matches that produced by the model (blue line) in response to the same paradigm. Simulating a flocculus lesion in the model (purple line) by removing the four forward models produces a complete abolishment of adaptation, whereas an NPH lesion (orange line) left the adaptation intact.

### Effects of lesions

In the model we simulated a lesion of the flocculus and a lesion of the NPH. The way in which this should be done in the model depends on the role that is ascribed to either structure (see Discussion).

### Flocculus lesions

We modeled a lesion of the flocculus by removing all the Forward Model boxes (Hexagon boxes in Figure 1). Figure 11A shows the result. The OKR is virtually absent. Meanwhile VOR gain is increased, and VOR phase increases at low frequencies. Following a model floccular lesion, the VOR did not adapt (Fig 10).

**Figure 11.**
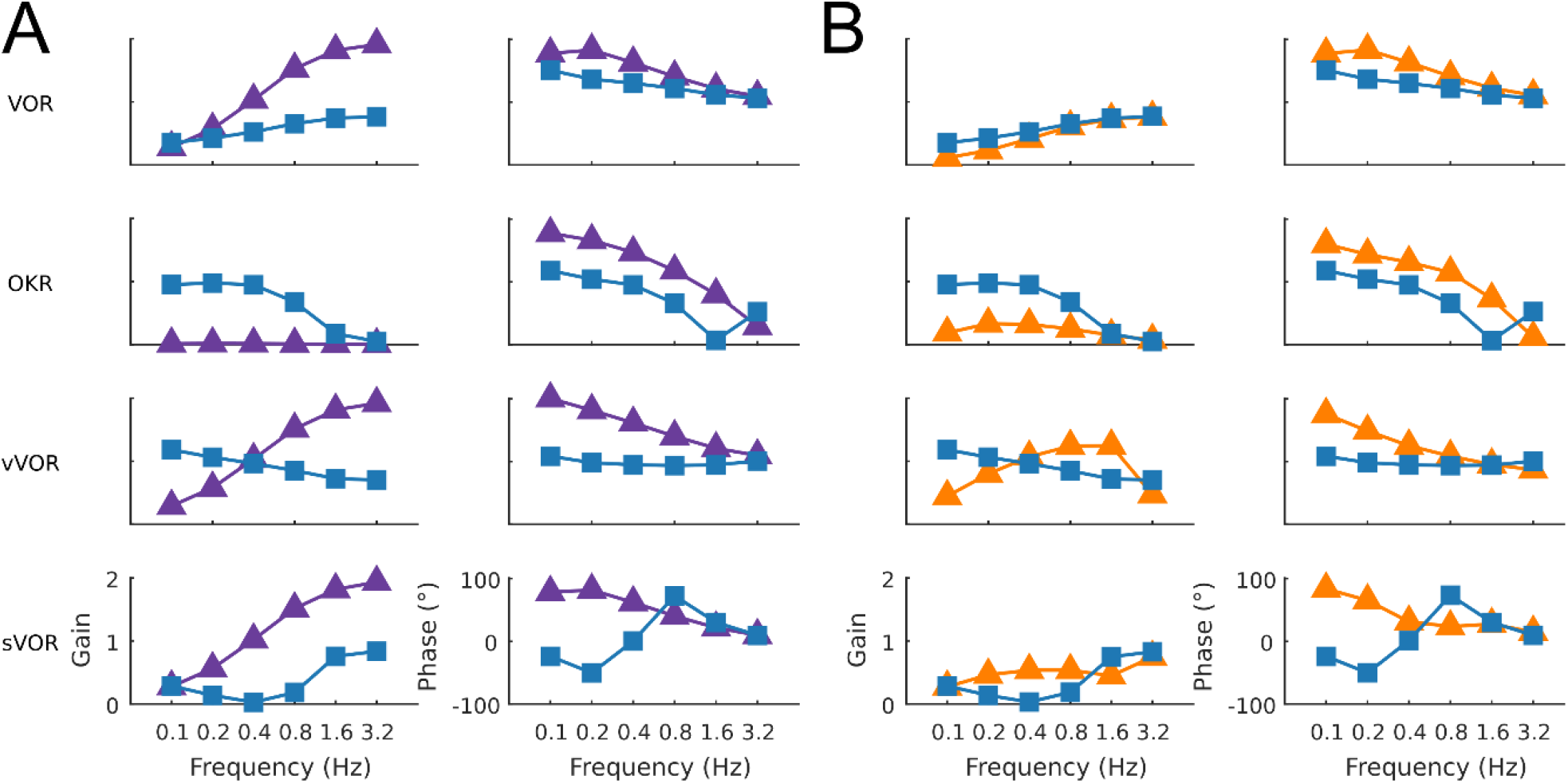
The effect of simulated lesions of the flocculus (A) and NPH (B) in the model on compensatory eye movements. The intact (blue line) and lesioned model response are summarized in Bode plots for the four conditions with the gain and phase presented in the left and right columns respectively. Following a simulated flocculus lesion removal of the forward model stage produces an increase in the VOR gain and phase and an almost complete loss of the OKR response. Due to the loss of the OKR component the response in the vVOR and sVOR conditions is almost identical. Similarly, the greatest effect of a lesion of the NPH was on the OKR response with a large decrease in gain and decrease in phase lag

### NPH lesions

If one believes the NPH to be part of the controller (Green et al., 2007), a lesion of the NPH would mean removing the inputs of the two outer hexagonal Forward Model boxes of Figure 1. A lesion of the flocculus would then be setting the values of all Forward Model boxes to a constant value of 0.

Alternatively, if one believes the NPH is the oculomotor integrator (Cannon and Robinson, 1987), an NPH lesion means setting the output of (outer, hexagonal [Fig 1]) integration boxes to 0. A lesion then only affects the two inner FM boxes of Figure 1 (“post-VOR slip” and “uncompensated slip”). We tested both manipulations.

Both types of lesion of the NPH resulted in exactly the same result. This is not surprising, since they are equivalent to setting the input to the integration step to 0, or setting the output to 0. Both produced a small effect on the VOR with a decrease in gain at low frequencies, reflecting the mainly feed forward nature of response. OKR in contrast was greatly affected with a large decrease in gain (Figure 11B). As expected (see discussion), the lesion also had an effect on the drift of the eyes back to the center in the dark, decreasing the time constant from 2.83s to 0.31s. Stahl et al (2006) report a time constant on the order of 5s for the neural integrator in C57BL/6 mice, although there was considerable variation between mice and over time.

Cheron et al (1986a, 1986b) made lesions in the NPH of cats. They show that such a lesion reduces low frequency VOR responses and completely abolishes OKR. However, the gain and phase measurements do not depict the full nature of the changes in the response to OKR. When applying low velocity stimuli, the OKR in our model becomes noisy and dominated by oscillations at the time points in which stimulus velocity is highest (Fig 12).

**Figure 12.**
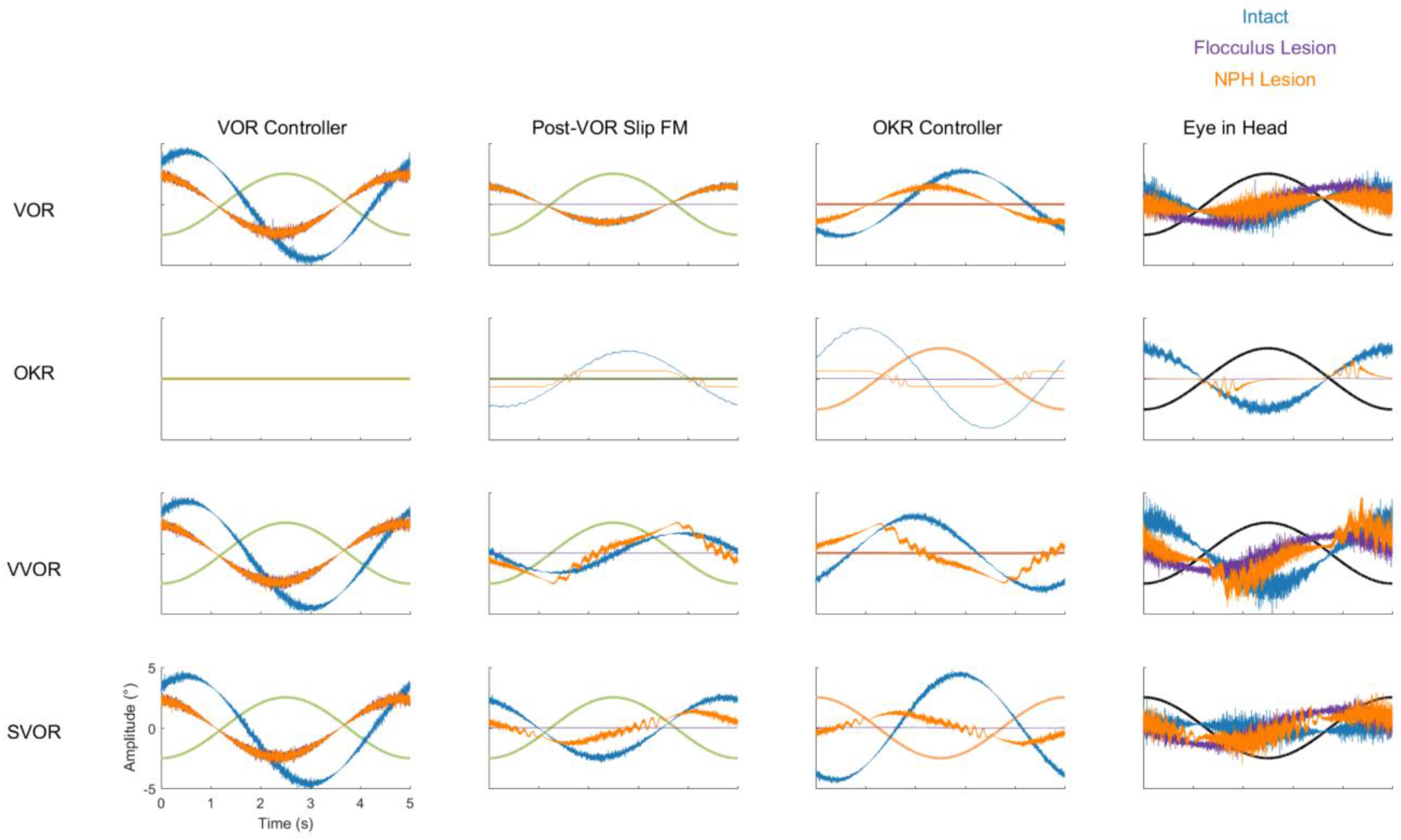
Summary of the dynamics of key components of the model in all four stimulation conditions (VOR, OKR, vVOR and sVOR) for the full model (blue) and the simulated Flocculus (purple) and NPH lesions (orange). The first column of plots represents the output of the VOR controller, second column displays the forward model of Post-VOR Slip, the third and fourth columns depict the commands produced by the OKR controller and the Plant respectively. In all plots the relevant stimulus is also displayed.

In our model, NPH lesions do not affect adaptation to the sVOR stimulation at 1Hz (Fig 10), because the individual reflexes at that frequency are relatively unaffected, and the site of plasticity is not lesioned.

### Effect of Lesions on Dynamics

To better understand how the different lesions affect the internal dynamics of the model, Figure 12 presents the post-VOR slip and the activity of the visual, vestibular and combined controllers in each of the lesion conditions for each of the four different stimulus conditions. There are a number of key findings. First, both the floccular and NPH lesion have the same effect on the vestibular command. This is because both lesions impact the vestibular command by eliminating forward model estimation of eye eccentricity. This leads to a decreased amplitude and increased phase lag in the vestibular command. Meanwhile, it can be clearly seen that the NPH lesion primarily affects the magnitude of the OKR.

## Discussion

### Brief summary of results

Frens and Donchin (2009) proposed that CEM can be modeled by an SPFC framework where specific functional roles can be ascribed to specific nuclei in the CEM circuitry. Here, we measured –for the first time-VOR, OKR, vVOR and sVOR over a large range of frequencies and amplitudes in the same animals. We then implement the SPFC framework in a detailed computational model which can, with a single set of parameters, mimic the behavior of OKR and VOR (Fig 3, 4 and 7). With the same set of parameters, the model also reproduces vVOR, sVOR (Fig 5, 6 and 7) and non-periodic SoS-stimuli (Fig 9). Furthermore, it successfully predicts the effects of lesions (Fig 11 and 12) and has adaptive behavior, similar to VOR learning (Fig 10).

The strength of this model is that it has relatively few critical parameters (see table 1) and that the critical parameters can be straightforwardly experimentally derived. This is an advantage over other SPFC-like models that address other motor systems (Shadmehr and Krakauer, 2008).

Key to the model are two distinct circuits for VOR and OKR. The VOR loop is relatively simple, and mainly consists of an integration step. In traditional models (for review see Glasauer, 2007), the OKR responds to actual retinal slip. However, due to the relatively long delay of the visual processing, the OKR response would then typically respond late. OKR state estimation in our model resolves this by predicting retinal slip. Both the VOR and the OKR loop contribute to this internal estimate of (uncompensated) retinal slip. This combined contribution is necessary, since the OKR assumes that the vestibular system will only partially resolve the retinal slip. While the reality may be more complex, the idea that the OKR models the VOR was the only way that we could explain the relatively high gains of both the OKR and VOR systems in isolation with the veridical gain of the two systems combined.

Finally, our model implements adaptation as a recalibration of this OKR estimate of VOR slip compensation. This helps explain why floccular lesions have a stronger direct effect on OKR but also disrupt VOR adaptation.

### The non-linear response to SoS stimulation

In addition to reproducing the response to sinusoidal stimulation in a wide range of conditions, the model also matched responses to SoS-stimuli that are identical to those previously used by (Sibindi et al., 2016). Strikingly, two non-linearities reported in the results of that study were reproduced: The first is that when confronted with a stimulus that consists of two non-harmonic optokinetic sinusoids, the amplitude of the lower frequency is suppressed, independent of the absolute value of the constituent frequencies. This then also results in changes in the amplitudes in vVOR and sVOR conditions. The second is that the lag of the response to the lower frequency is larger, resulting in a delayed overall response. This can be seen for both VOR, OKR and its combinations.

The model has one non-linearity specifically built in: the saturation of the visual motion sensitive neurons in the retina (see Eq 8 in the Supplementary Material). Explicitly removing this saturation eliminated the gain decrease and delay increase of the OKR and vVOR, but left the increased delays in the VOR and sVOR unaffected (Fig 9).

These modeling results support the hypothesis that Sibindi et al. used to explain their results: increased delays may be a result of the circuit properties. That is, they suggest the forward model fails to predict upcoming retinal slip in complex stimuli. Our results also support their hypothesis that the gain changes are probably the result of non-linear retinal processing.

### The role of the flocculus

The flocculus acts as a forward model for both the VOR and the OKR loop. However, the role it plays in each reflex is completely different. The flocculus is not critical for VOR performance, as animals lacking Purkinje cells do have an intact VOR although the amplitude of the response is significantly higher (van Alphen et al., 2001). While our model does include a forward model and state estimator for head velocity, this is only a formal result of the structure of the model. In fact, our model ignores the results of the forward model and uses the sensory information exclusively to determine head velocity. Thus, the role of the forward model (green hexagon in Fig 1) in this system is actually only to integrate eye velocity into eye position. For the OKR loop the forward model helps to overcome the delay in the OKR feedback loop, and it is crucial to provide information about the estimated post-VOR slip.

We mimicked lesioning the flocculus by removing the output of the forward models. This removed the capability of the system to predict upcoming retinal slip. As a result, the optokinetic response was virtually abolished whereas VOR gain substantially increased (Fig 11A). Lurcher-mice, a mutant strain that lacks Purkinje cells, have substantially lower OKR gains than their wild type littermates (van Alphen et al., 2002). Lurcher-mice results are also similar to a floccular lesion in our model in that VOR-gain is increased. Results on VOR gain in acute, non-genetic floccular lesions are mixed (Rambold et al., 2002).

We can understand the results showing increased VOR gain in Lurcher mice using our model: the OKR generally acts to suppress the VOR and a floccular lesion releases this suppression. This interpretation leads to the further prediction that floccular lesions will reduce the effect of visual suppression of the VOR, increasing gains in the sVOR. This is true in our model as well as being compatible with the literature (Belton and McCrea, 2000; Takemori and Cohen, 1974; Zee et al., 1981).

The change in phase of VOR response that is seen in Lurcher mice (van Alphen et al., 2002) can be modeled only if we include the VOR integration stage in the flocculus. This supports the view of Green et al (2007) that the NPH provides an efference copy that is integrated in the flocculus (see below).

### The role of the NPH

Our model provides a potential resolution to a debate about the role of the NPH in eye movement generation. In Robinson’s inverse-model framework, the NPH is thought to act as the neural integrator for horizontal eye position. Such an integrator is necessary to provide the abducens nucleus with both velocity and position commands that are needed to overcome the low-pass filtering properties of the plant (Robinson, 1981). This view has been widely adopted by researchers in the oculomotor system. A critical finding supporting this view is from Cannon and Robinson (1987) showing that lesions of the NPH cause the eye to drift towards the center of the oculomotor range. This is compatible with the loss of an integrator that opposes the elastic restoring forces of the plant. However, more recently Green et al. (2007) showed that the burst tonic neurons of the NPH have activity that is nearly identical to that of the motor neurons in the abducens nucleus. Furthermore, these neurons have direct projections to the flocculus (Belknap and McCrea, 1988; Langer et al., 1985; McCrea and Baker, 1985). On the basis of these findings, they proposed that the NPH provides efference copy input to a cerebellar forward model (Ghasia et al., 2008; Green et al., 2007). This view was also incorporated in our SPFC (Frens and Donchin, 2009). Thus, in our model, an NPH lesion removes input to the forward models. However, when we lesion the NPH projection in our simulation (by removing efferent copy to the forward model or by removing its output), we found that we had reproduced the Cannon and Robinson (1987) result: the time constant of the drift was reduced. Hence, a lesion of the efference copy projection produces the same results as those thought to support the idea that NPH is an integrator. It seems that the Canon and Robinson (1987) results are compatible with both models while recent anatomical and physiological findings support the idea of efferent copy.

### VOR Adaptation

Within our framework, VOR adaptation happens through adaptive changes in the forward model of VOR used by OKR. OKR assumes that VOR will correct a certain fraction of sensed head velocity. Determining the proportionality constant robustly led to the same value regardless of stimulus amplitude over a wide range of frequencies. When challenged with an adaptation stimulus, the model gradually changed its gain. Of course, the rate of adaptation could be set arbitrarily. Our setting led to an adaptation speed that is very similar to what we experimentally found in mice under identical experimental conditions. To our knowledge, we are the first to suggest that VOR adaptation reflects adaptation of a forward model of VOR output. However, the idea is compatible with the recent suggestion that VOR adaptation is driven by the motor consequence of retinal slip rather than the slip itself (Shin et al., 2014). Floccular lesions in our model abolish VOR adaptation, which is in line with the literature (Schonewille et al., 2010). NPH lesions do not affect adaptation at 1 Hz in our model, but to the best of our knowledge there is no literature to corroborate this finding.

Although our model is capable of adaptation, we believe that adaptation in the biological system is probably more complex than that in our model. Biological adaptation seems to reflect plasticity at multiple sites with multiple time constants (Clopath et al., 2014; Gao et al., 2012; Porrill and Dean, 2007). The introduction of more realistic adaptation and testing adaptation at higher and lower frequencies is an important future extension of the current model.

### Relationship to Other Models

The CEM system is a popular candidate for computational modelling due to the known anatomical substrates and the restricted degrees of freedom. Theories of motor control are primarily based on one of two main architectures. One theory suggests that the motor system relies on generating an ideal “desired movement” or “desired trajectory” that serves as a basis for subsequent control. Such an architecture faces a number of key challenges: generating the desired trajectory, translating it into motor commands, and correcting for deviations during online control. At the heart of such a system is an “inverse model” which translates desired movement into motor commands (Jordan and Rumelhart, 1992). The literature in the CEM system contains a long tradition of such models (for example: Clopath et al., 2014; Glasauer, 2007; Kawato and Gomi, 1992; Lisberger, 2009; Robinson, 1981). In general, a desired motor command is fed to the brainstem, which then acts as an ‘inverse plant’, i.e. it processes the command in order to overcome the low-pass properties of the extraocular muscles and tissues that are connected to the eye.

The key innovation in our model is the use of recurrent cerebellar-vestibular nuclei loops which enable the model to function correctly in the presence of considerable motor and sensory noise and in the presence of significant delays in sensory feedback. There exists anatomical evidence for such loops (Büttner-Ennever and Büttner, 1992) and proposals for their functional significance have been made previously (Porrill et al., 2004).

Since the optimal control framework was originally proposed as an approach to understanding vertebrate motor systems, models of this sort have been implemented in the control of various motor tasks. The implementations closest to our model are those that attempt to describe coordinated head and eye movements during gaze shifts (Sağlam et al., 2011, 2014; Todorov and Jordan, 2002). One somewhat similar model has been proposed to describe the CEM system (Haith and Vijayakumar, 2007).The Haith model is built largely to address adaptation to changing dynamics, an issue not addressed by our data or our model. Additionally, the Haith model is not confronted with actual data. In sum, our model is unique in a number of respects: (1) the extensive data with which it is challenged, including lesion data and non-sinusoidal data, (2) the idea that one of the main drivers of adaptation is compensation of the OKR system for predicted VOR error, (3) the development of a fully realized recurrent model of the CEM system in the spirit of the optimal control feedback framework.

## Acknowledgements

This research was supported by the C7 Marie Curie ITN initiative (TS, PH, SD), TC2N Interreg Grant (OD, MF), the ABC Robotics Initiative (OD) and a Post-Doctoral Fellowship from the Kreitman School for Advanced Studies at BGU (PH).

## Supplementary Material

### 1 Overview of Model

This section describes the details of the model of the CEM described in the main text. The description provides all of the equations used in sufficient detail for the model to be implemented, although the actual Matlab code is available on the Open Science Framework website (https://osf.io/feq7c/). The model was implemented in Matlab R2016a (The MathWorks, Natick, MA). The time step for the simulation used was 1 ms.

This section is divided into subsections that describe the implementation of the plant and the control system. In the section on the plant, we describe both the effector and input implementations. The effector implementation is a model of how firing in the oculomotor nuclei affects muscle activation, and how that drives eye movement. The inputs we model are the vestibular and the retinal inputs to the system. The description of the control system is divided into three parts: the actual state dynamics; the system’s estimate of state; and the transformation of state estimate into motor command.

### 1.1 The plant

In this section we describe the dynamics of eye movement as a function of the firing rate of neurons in motor nuclei (OMN/AB) that project to eye muscles. Output of the OMN/AB innervates the horizontal rectus muscles, which are responsible for horizontal eye movements. These nuclei are reciprocally activated and project to muscles that move the eyes in opposite directions. Hence eye velocity depends on the difference between OMN and AB activities. The transfer function of these nuclei for the monkey has been described using the formula (Robinson, 1981):

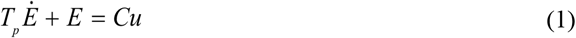

(Where *E* is eye position, *u* is motor command from the OMN/AB, and *C* and *T_p_* are the gain and time constants, respectively). The motor commands from the two nuclei were not separately modelled, but rather their activity was represented in a combined manner as the sum of two oppositely signed command signals.

Eq. (1) describes a leaky integrator with leakage time (in s). In monkey, *T_p_*has been estimated at 0.24s and in rabbits it can be estimated from the work of Stahl and Simpson (1995) and more recently for mice in Stahl et al. (2015) to be 0.5s. We ran our simulation both with *T_p_* = 0.24 s and with *T_p_* = 0.5 s, and saw no difference in the results. For this paper, we present results using *T_p_* = 0.5s (see Table 1). For the purpose of the model, we absorbed the constant *C* into the definition of *u*, so that our motor command was specified in °/s rather than in units of firing rate:

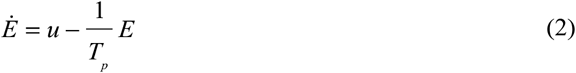

### 1.2 Sensory Signals

Compensatory eye movements are driven by two different sensory signals – vestibular and retinal. In this section we describe the biological processes behind these sensory signals and the numerical models that can be used to describe them.

### 1.3 Vestibular input

Vestibular input is created by the semicircular canals in the inner ear. We transformed the head velocity to sensory signal in three steps: linear filtering, velocity-sensitive transformation, and delay. At high frequencies, canals sense head rotation velocity with high accuracy. However due to the physical properties of the sensor, the accuracy is not good at low frequencies

(Robinson, 1981). Thus, the semicircular canals can be best described as a high pass filter that acts on head velocity:

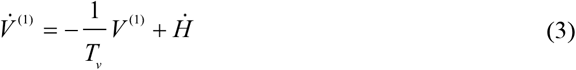

Where *V* ^(1)^ is the first stage of the neural signal generated by the velocity sensitive vestibular afferents (as opposed to *V*, the internal representation of head velocity) that are driven by the actual rotational head velocity, *Ḣ*, and *T_V_* is the filter constant that defines the effective sensitivity range of the afferents. The value of *T_V_* differs between species. In mice this constant was measured in Yang and Hullar (2007). While they fit their data using a fairly complex transfer function (here reproduced in its original Laplace-domain notation):

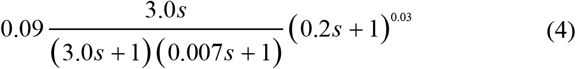

A first order approximation of the formula, and neglecting the leading constant, gives us Eq. (3). Over the relevant frequency range, the two functions are nearly identical, with *T_v_* = 3sec for regular afferents of the horizontal semi-circular canal that project to the vestibular nucleus. Van Alphen et al. (2001) found that a lower time constant is needed to explain VOR experimental data. It is possible that additional filtering in the input synapses of the vestibular nucleus explains the difference between the constant measured in the afferents and that seen behaviorally. However. we found that our behavioral data was best matched by a constant very close to that found by Yang and Hullar (2007), *T_v_* = 4 sec.

Subsequently, we introduced a delay and added noise:

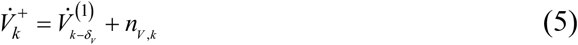

The vestibular delay (δ*_V_* = 2 ms) represents the physical response time of the semi-circular canal and the neuronal transmission delay (Sohmer et al., 1999). The noise (*n_V_* _,*k*_) has a standard deviation proportional to the size of the vestibular signal (with constant of proportionality *a_V_*, with the tilde,∼, meaning “distributes as” and *N*(*µ, σ*^2^) is the normal distribution with mean *µ* and variance *σ*^2^):

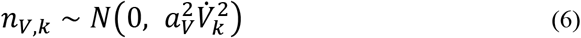

Since vestibular inputs depend only on head movement and head movement is determined by the experiment, the behavior of the system has no effect on vestibular inputs. Thus, we calculated these signals offline before running the simulations and introduced them directly as input.

#### 1.3.1 Retinal Input

Visual information is provided by motion sensitive neurons in the retina (Yoshida et al., 2001). Those neurons sense local velocity of the image on the retina (often called retinal slip). In our experiments, the entire retina experiences the same retinal slip, and it is equal to:

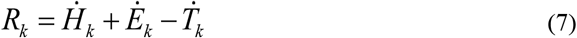

Where *R* is retinal slip velocity, in °/s, *Ṫ* the velocity of the visual surroundings in °/s, and *Ė* is the velocity of the eye relative to the head (generated as described above in Eq. (2).

The retinal motion sensitive neurons are linear in a limited range. In rabbits, sensitivity peaks at about 0.6 °/s (Oyster et al., 1972), with neuronal firing rates increasing through this range, but then dropping off for higher velocities. At 10 °/s the neurons are unresponsive. Neurons in the AOS (the retinal target driving OKR) have shown similar properties (Soodak and Simpson, 1988). Currently available data does not give the precise saturation point for the motion processing system of the mouse. In order to fit our data, our model assumes saturation of *R*_max_ = 0.65 deg/sec and a piece-wise linear response function, representing a population code of neurons that individually drop off at values between 0 and R_max_:

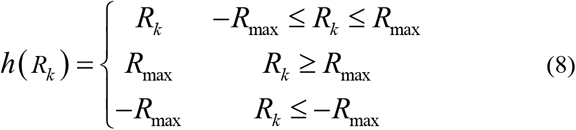

The processing of visual signals adds substantial delay to the retinal feedback (Collewijn, 1969). Our model uses the value of *δ_R_* = 70 ms proposed for the delay in mice (van Alphen et al., 2001):

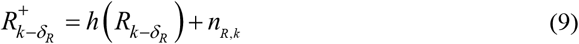

With 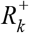 the current internal representation of retinal slip, and *n_R_* _,*k*_ being the retinal noise, which has standard deviation proportional to the retinal activation (with a constant of proportionality 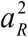):

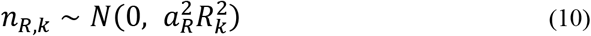

#### 1.3.2 Full system dynamics

The above descriptions of the oculomotor plant and the retinal and vestibular input are combined to make a nearly linear state equation for the plant. Thus, we use a standard linear systems formulation (Frens and Donchin, 2009) with the state of the system at time *k*, *x_k_*, undergoing a particular dynamics specified by the matrix *A*. In addition, the state is influenced by three factors: the command signal, *u_k_*, affects the state through a matrix, *B*, that specifies how each part of the command signal influences each element of the state; the external input, *z_k_*, represents the influence of the external world on the state; also, the state is influenced by noise, *n_k_*. Finally, this state leads to sensory input (often called the observation), *y_k_*, through a matrix, *D*. Altogether, this leads to what is called the system equations:

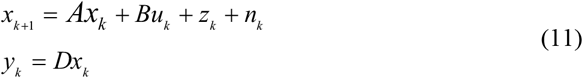

These system equations are linear. Each piece of this equation is treated in detail in the paragraphs that follow.

The state at time step k is represented by the following vector:

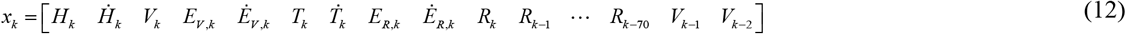

The state includes time-delayed versions of the retinal and vestibular sensory signals. *R_k_* represents the retinal input being generated at this instant (based on the current eye velocity) and *R*_*k*−1_ through *R*_*k* − 70_ represent increasingly delayed versions. The observation matrix, Eq. (20), is such that only the fully delayed retinal slip, *R_k_*_−70_, is available to the state estimation. The vestibular input is not affected by the behavior of the system, so it was generated offline according to Eq. (3) and delayed by 2 ms according to Eq. (5).

*z_k_* is the external input and includes the change in the actual head velocity, vestibular sensory signal, and movement of the visual stimulus. These signals can all be generated offline before running the simulation. The vector can be written as:

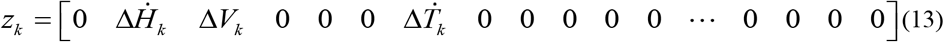

*n_k_* is the noise in the system. It affects eye velocity as well as vestibular and retinal input, so it can be written as:

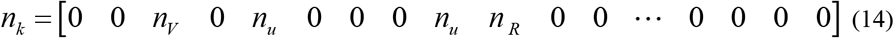

In modelling the noise, we opted for model simplicity over realistic modelling of the noise. We followed the general idea in Todorov (2004) and Harris and Wolpert (1998) of having the size of the noise be proportional to the signal. Vestibular noise and retinal noise have already been described in Eqs. (6) and (10) respectively. The standard deviation of the motor noise is similarly proportional to the motor command (with constant of proportionality *a_u_*)

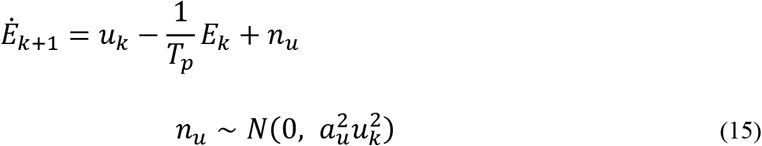

We ran the model with different constants of proportionality for the noise (*a_u_*, *a_R_* and *a_V_*) up to 0.5 and did not see a change in the results. Given that we have no available data on amount of sensory or motor noise in the system we used values well in the middle of stable range, i.e.:

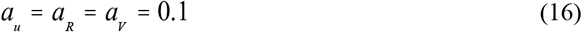

*A* is the matrix describing the state dynamics and is written as:

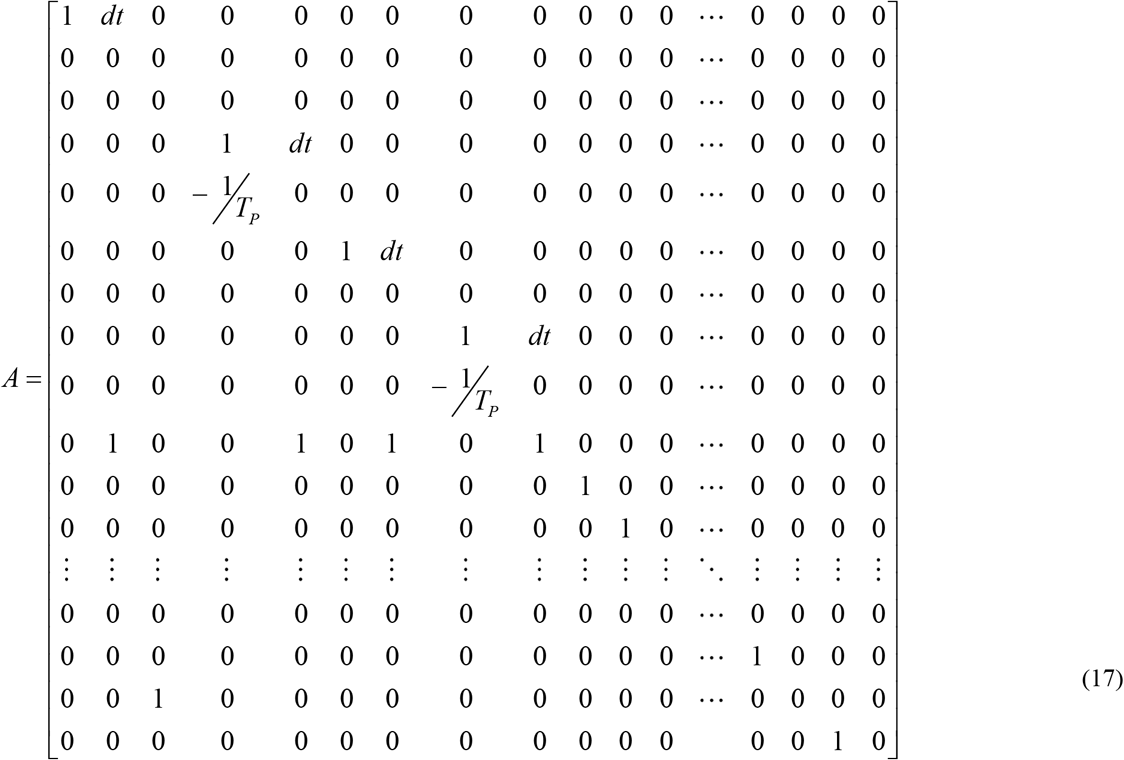

Rows 2, 3 and 7 (velocity of the surroundings and of the head and the vestibular signal) are all just equal to 0. This reflects the fact that these variables are controlled by the inputs and not part of the dynamics of the system, in our model. Row 4 (eye position) simply includes the change in eye position caused by eye velocity (column 5), which needs to be scaled by *d =* 0.001 because eye velocity is in units of °/s and the time step is 1 millisecond. It is worth noting that row 8 also describes eye dynamics (just like row 4). These representations are separated because in the internal controller they reflect different estimates. The simulation code keeps them in register by replacing them with the sum of the two values on each time step. Row 5 (and row 9) describe the tendency of the eye to drift back to center (the position dependent part of Eq. (2)). Row 10 says that current retinal slip is equal to head velocity plus eye velocity minus stimulus velocity (Eq. 7). The rest of the dynamics matrix (rows 13 through 78, not shown) simply shifts previous measured retinal input backwards in time (e.g. *R*_k_ → *R*_k−1_, *R*_k−1_ → *R*_k−2_).

State transition is not, however, strictly linear. This non-linearity is represented by the function *h*(*Ax_k_*) in Eq. (8) so that,

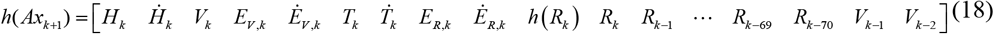

Where *h*(*R*) describes the saturation of the retinal sensory signal (Eq. (8)). That is, every element of the state vector is preserved by *h* except the retinal slip which saturates.

Since the motor command, *u_k_*, is a scalar, the control matrix B of Eq. (11) is a vector with the same size as the state. Because the command affects eye velocity directly, the only non-zero element of B is in the row representing eye velocity. Units are adjusted so that 1 unit of motor command (neural activation) causes an acceleration of 1 °/ms, so *B* is:

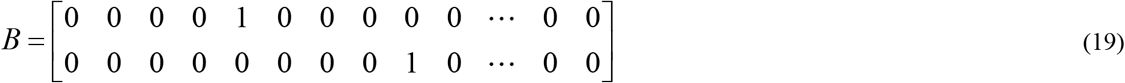

The second equation in Eq. (11) describes the observation, which is the part of the state available to the controller. The observation vector, *y_k_*, contains delayed retinal and vestibular inputs. Thus, it can be calculated linearly using the observation matrix *D* (which is simply a 2×82 matrix of zeros with ones at locations (1, 82) and (2, 80) for vestibular and retinal input respectively). The *D* matrix is applied to the retinal slip after saturation, and we also add in sensory noise at this stage.

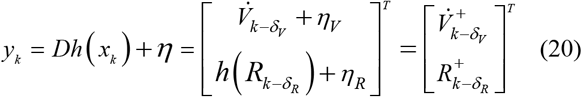

#### 1.3.3 Control system

In this section we describe an optimal feedback controller for the compensatory eye movement system. This controller includes a forward model and a process of combining forward model prediction with sensory input, called state estimation. We will use the hat notation, 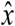, for estimates produced by the forward model and the tilde notation, 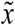, for the combined state estimate.

The operation of the controller can be described globally with the following equations:

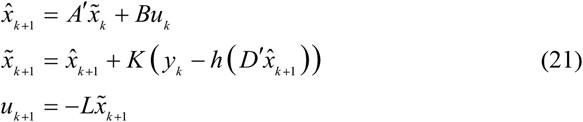

The first equation says that the forward model uses the previous state estimate and the previous motor command to generate a prediction of the next state. The second equation says that the estimate of the next state is generated by correcting this prediction for discrepancies between predicted and experienced retinal slip. The last equation says that motor command will be a linear function of the state. The tags on some symbols result from the fact that the controller’s internal representation of state is different from the actual system state. Thus, *A*′ is the internal representation of system dynamics and *D*′ selects the appropriate sensory inputs from the internal system state.

#### 1.3.4 VOR control

Our model assumes, as described in the main text, that VOR and OKR involve separate neural processing. Thus, it will be clearest if the operation of each is described separately, and then the combined matrix equations will be easier to follow.

The architecture of the VOR is the same as the overall architecture of the system:

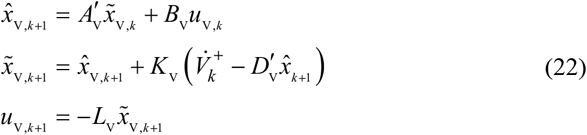

In the case of VOR, since we have no access to the actual head velocity, we use the vestibular signal as an approximation of the head velocity. Thus, the state needs only have five elements:

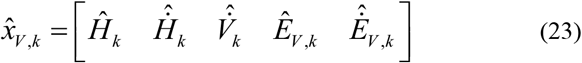

The forward model is quite simple. The head velocity is not affected by either system dynamics or command (row 1 of Eq. 24 and the first 0 in Eq. 25). Eye movements have the usual plant dynamics (rows 4 and 5, which are taken from Eq. 2) and are affected directly by the motor command (the 1 in the third fifth position of Eq. 25):

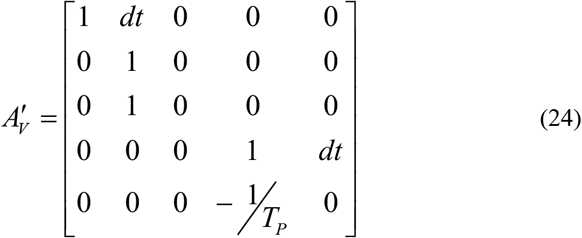

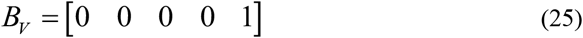

The observation matrix returns the estimated head velocity (which is what we expect the vestibular input to be):

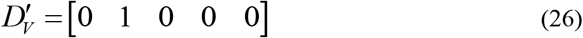

This is compared to the actual vestibular input, 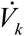. Because a floccular lesion does not eliminate VOR performance, we set *K_V_*=[0 1 1 0 0]. That is, the sensory feedback completely replaces the forward model in our knowledge of head velocity. The role of the forward model in this system is actually to integrate eye velocity into eye position.

Finally, the actual motor command is generated (see below for how these values are determined) using the equation 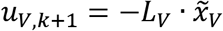 with 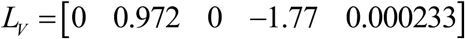 so that, ultimately:

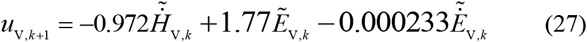

#### 1.3.5 OKR control

The job of the second part of the control loop is to estimate uncompensated retinal slip and compensate for it. Uncompensated visual slip arises from three sources: changes in the velocity of the visual stimulus, noise in the system, and head movements not compensated by the VOR. Importantly, the system cannot distinguish changes in the velocity of the visual stimulus from noise in the system. We use the symbol 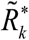 for the system’s estimate of all three of these quantities together: the retinal slip uncorrected by VOR. We also call this the post-VOR slip, and it represents how much the visual environment would be moving in the absence of OKR.

The OKR’s prediction of uncompensated retinal slip is thus the difference of two quantities: the post-VOR slip and the estimate of how much the OKR is moving the eye, 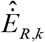:

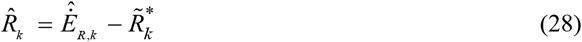

The OKR system assumes that some amount of head movement will be compensated for by the VOR. Its estimate of uncompensated visual input generated by sensed head velocity is proportional to the actual sensed head velocity. Our forward model estimate of uncompensated post-VOR retinal slip will be different from our previous estimate because it is updated by a factor proportional to head acceleration:

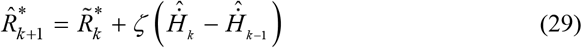

(where *ζ* is the constant of proportionality and is discussed in the section on VOR adaptation below). We then use a Kalman filter to incorporate sensory prediction error and produce a final estimate of post-VOR retinal slip:

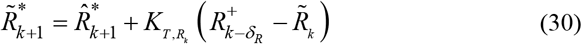

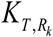 represents the appropriate term in the Kalman gain matrix (specified fully below). Our data was best fit by using *ζ* = −0.6 and 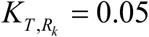 which means that that OKR has a tendency to overcompensate for head rotation and that it estimates that 5% of unexpected retinal slip represents real movement of the visual surroundings. Note that Eq. (30) also uses 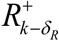 and 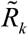 which are the currently available retinal slip and its estimate while Eq. (28) and Eq (29) used 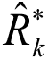 which is the estimate of the retinal slip happening right now. This estimate will be delayed for 70 ms before it becomes available as 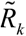. For the ease of the reader a supplement to Figure 1 (Figure S1-figure supplement 1) includes a version of the model schematic with the various forms of retinal slip and its estimates labelled.

With this understanding in place, we can describe the OKR control system. It has the same overall architecture as the full system:

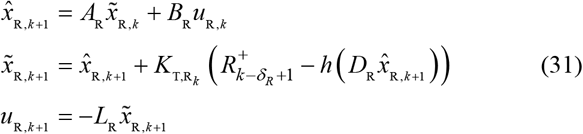

With function *h* representing the saturation of the retinal input (Eq. (8)). The state vector includes everything needed to calculate retinal slip, movement of the visual world, and head acceleration:

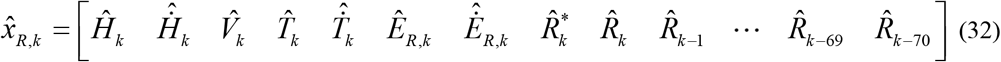

The forward dynamics matrix, 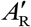, look like this:

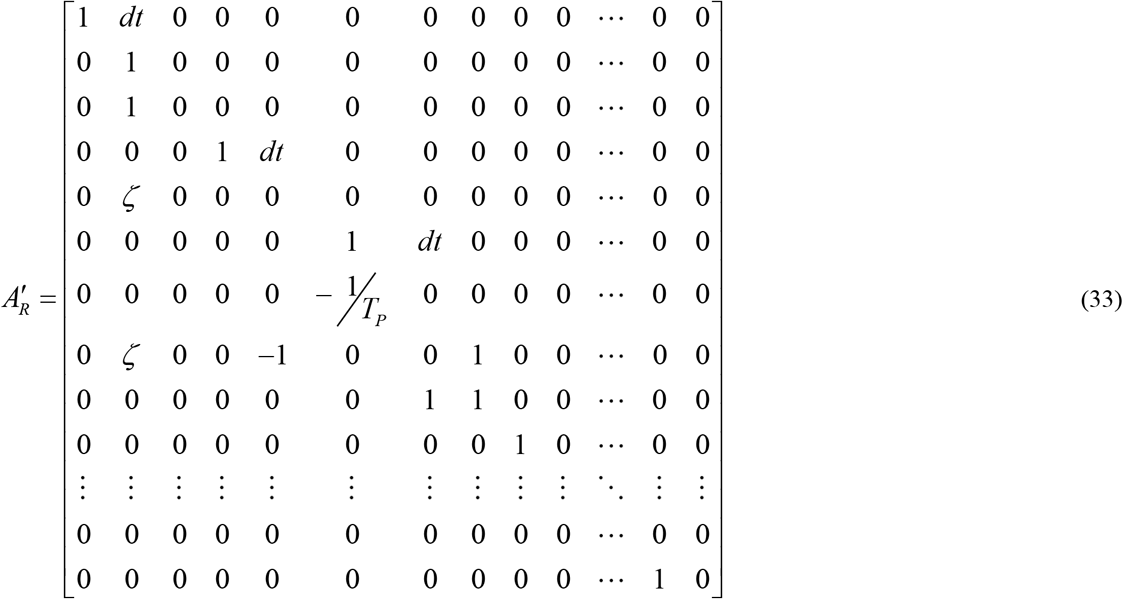

These rows accomplish: calculation of uncompensated post-VOR retinal slip (row 8, implementing Eq. (29)), shifting of current vestibular input to previous vestibular input (rows 4-5), modelling of the eye plant (rows 6-7, implementing Eq. (2)), calculation of the current uncompensated retinal slip (row 9, implementing Eq. (28)). The rest of the 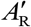 matrix takes care of the delay of the estimated retinal slip. The *B*_R_ matrix simply copies the motor command into the eye velocity vector, just as with the VOR system:

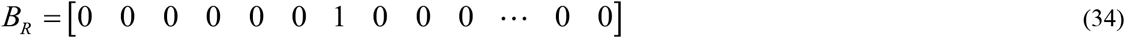

In calculating state estimation for the OKR system, we must take into account the non-linearity of the retinal processing before comparing the predicted retinal slip to the sensory input. We first use the matrix *D*_R_ to select only the predicted uncompensated retinal slip,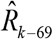, from the state vector, as in Eq. (26) but with a larger state vector. Then, the predicted uncompensated retinal slip is cut off with the saturation function of the retinal input, as specified in Eq. (8). This can be compared to the true retinal input 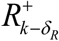, providing retinal slip prediction error. The retinal slip prediction error updates the estimated state values of post-VOR retinal slip and uncompensated retinal slip. Our data was best fit by using:

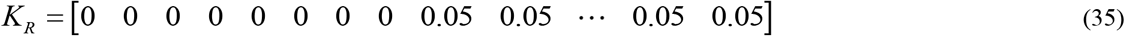

Finally, the motor command is generated by using the equation 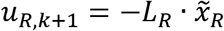 just like in the case of VOR (again, see below for derivations), with 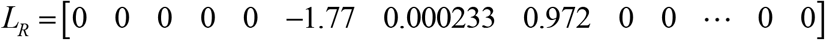 so that the motor command is:

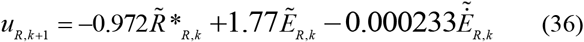

#### 1.3.6 The combined controller: forward model

To produce a combined system, as described in Eqs. (21), in our calculations we simply combine the descriptions of the OKR and VOR systems above. The only state variable that overlaps in the two systems is the head velocity. However, this poses no difficulties.

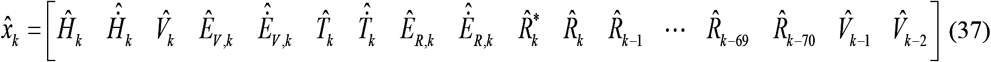

And the dynamics and command matrixes can be copied from the two systems described above (the last sets of rows just shift the retinal slip and vestibular input back in time):

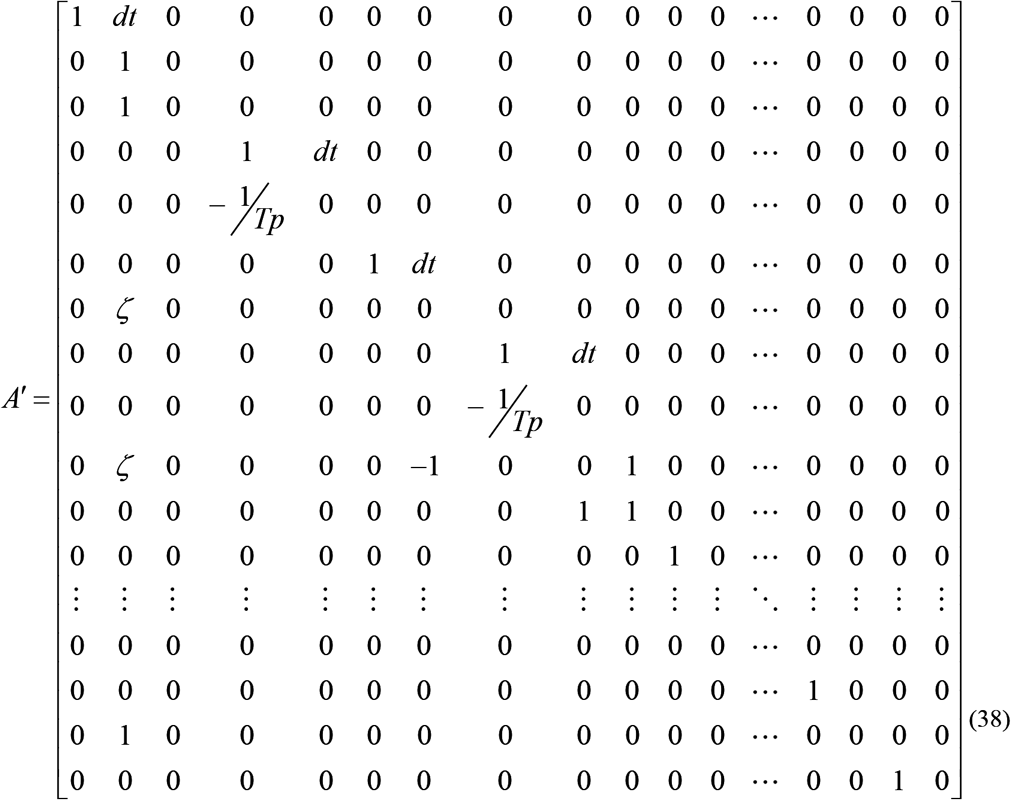

The internal representation of the command is two dimensional, with separate command for the VOR (dimension 1) and OKR (dimension 2), and each is added into the appropriate eye velocity:

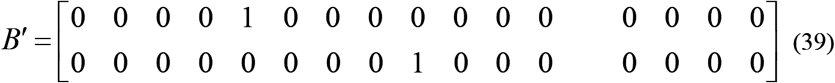

#### 1.3.7 The combined controller: state estimation

In the second equation of the set in Eq. (21), the observation matrix, *D′*, selects the vestibular and retinal input appropriately:

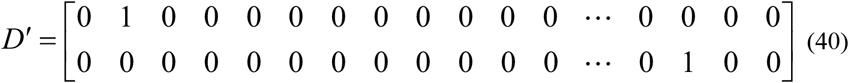

Note that the first row of *D′* is different than the first row of *D*. This difference comes from the fact that the internal system maintains an ongoing estimate of head velocity that is influenced by the input while the real system does not maintain such an ongoing estimate. The only representation of the delayed head velocity is the actual delayed head velocity. *h′*(*x*) applies the retinal saturation non-linearity, *h*(*R*) from Eq. (8), to the retinal slip and does not change the vestibular input:

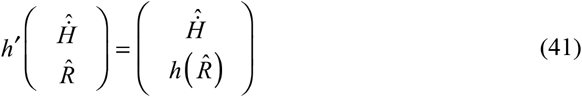

Parameters of the Kalman gain were selected by hand to match the data. We assumed that vestibular input only affects our estimate of the head velocity, 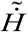, and that retinal input affects both our estimate of post-VOR retinal slip, 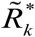, and our estimate of overall uncompensated retinal slip 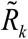 and its delayed versions. This gave the Kalman gain matrix the following form:

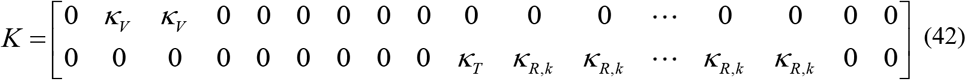

We set *κ_V_* to 1, in order match the experimental finding that floccular lesion does not eliminate VOR. We set the other values to match the behavioral data. That is, the larger the value of *κ_T_* and *κ_R_*_,*k*_, the more quickly new retinal input affects our estimates. When the Kalman gains for the visual system are too large, noise reverberates in the system, leading to an explosion of noise in the OKR at low frequencies. When they are too low, the system does not manage visual following. Balancing these two considerations, we got the best match for our data with *κ_T_* = *κ_R_*_,*k*_ = *κ_R_*_,69_ = *κ_R_*_,68_= … = *κ_R_*_,1_= *κ_R_*_,0_= 0.05

#### 1.3.8 The combined controller: cost function

We assumed that the primary goal of the optimal controller of the CEM in afoveate species (like rabbit and mouse) is to minimize motion of the visual field on the retina in order to stabilize the retinal image. We make the assumption that this cost is considered separately for VOR and OKR because we are assuming that these reflexes are supported by separate neural substrates.

Thus, the overall cost of the system can be broken down into two parts, vestibular and retinal:

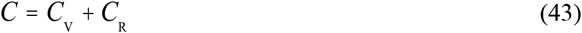

Each of the two sub costs is concerned with a different retinal slip: *C*_V_relates to 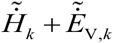, retinal slip due to uncompensated head motion, while *C*_R_ relates to 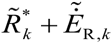, retinal slip due to uncompensated motion of the visual environment. In addition to the cost associated with retinal slip, each cost function includes a cost associated with eye eccentricities (this can be considered an “action” cost since eye eccentricity leads to extra muscle activity and energy expenditure). Finally, both cost functions discount future costs, as is common for an infinite horizon feedback controller: Thus, the two cost functions required for creating the two motor commands are:

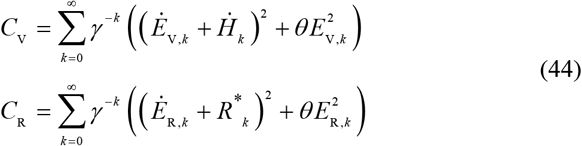

The parameter *θ* balances between eccentricity and retinal slip costs. The parameter *γ* is the discount parameter Bradtke (1993) used to reduce the influence of increasingly distant costs. These two parameters were needed to match the drift of the eyes in the dark and were set to *θ* = 2 and 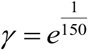. For simplicity we approximated the infinite sum in Eq. (44) with a finite sum; we kept the first 100,000 terms.

#### 1.3.9 The combined controller: the motor command

If our system had a linear plant (L), quadratic cost function (Q) and independent, identically distributed (i.i.d.) Gaussian noise, it would be called an LQR system (Åström and Murray, 2008). For such systems, it can be proven that the optimal controller can be separated in two independent parts – an observer and a simple controller – using the Ricatti equations (Lancaster and Rodman, 1995). We do not go into the details of these equations here, but we note that the CEM system, as described above, is not linear (because of non-linearities in the inputs) and does not have i.i.d. noise (since we use signal dependent noise). Nevertheless, the convenience of the LQR formulas has led to their frequent use in systems that are close to being LQR (Burns and Ou, 1994; Lopez-Martinez et al., 2004). Previous experience is that this leads to nearly optimal controllers, and we followed this strategy here.

However, before we apply Ricatti equations, we make one additional assumption. We assume that for the purposes of this solution, the controller assumes full correction of the head velocity by the VOR system. That is, we set *ζ* = 0 in the matrix *A′*, Eq. (38).

Applying the equations of Lancaster and Rodman (1995) to our system, Eq. (21), we derive a solution for the control policy, *L*.

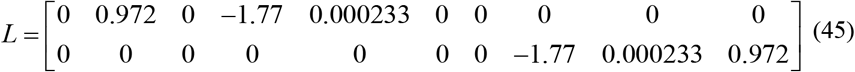

This can be more clearly written in terms of the final results for the motor commands:

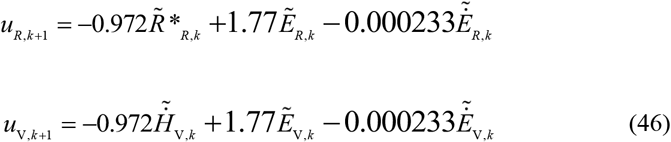

The first term in both Eqs (46) compensates for retinal slip. The second term combines compensation for the “drift to center” generated by the elastic properties of the plant (Eq.(2)). This activity is apparently generated by the “neural integrator” produced by the firing of the tonic and burst-tonic premotor cells (Robinson, 1981). Experimental results presented in this article and in other works (Cannon and Robinson, 1987) show the elastic properties of the plant are not fully compensated for by the controller; i.e. the neural integrator is leaky, and this leakage has a much higher time constant than the elastic term of the plant.

#### 1.3.10 VOR adaptation

The parameter ζ (introduced in Eq.(29)) represents the extent to which the OKR system assumes head movements will go uncompensated. We model CEM adaptation as adaptation of this parameter so as to accurately predict retinal slip. The forward model prediction of retinal slip is given by Eq. (28). Where we recall that the star indicates that this is the estimate of the retinal slip that is we predict that is happening right now (post-VOR slip), as opposed to the estimate of the available retinal slip (with a 70 ms delay) which is indicated by 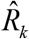.

We want to minimize the error in retinal slip prediction error (Figure 1-figure supplement 1):

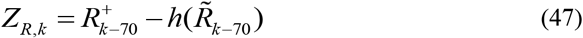

We employ a decorrelation approach to adaptation (Porrill et al., 2013) and update ζ based on a factor proportional to the correlation of head acceleration and retinal slip prediction error.

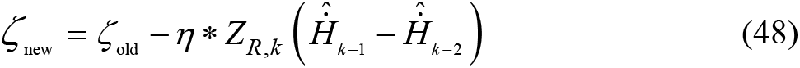

Where *η* specifies the rate of adaptation. For the results presented here ζ was updated every 4 cycles of the stimulus (although this value is not critical and adaptation functions correctly with a wide range of update schedules) and was set to match the rate of adaptation in the experimental data: *η =* 0.018.

## Experimental Methods

### 1.4 Animals

In order to test the model we recorded CEM in 13 C57Bl/6J mice (Charles River, Wilmington, MA, USA). All mice were housed on a 12h light / 12h dark cycle with unrestricted access to food and water. Experiments were performed during their light phase. All experiments were performed with approval of the local ethics committee and were in accordance with the European Communities Council Directive (86/609/EEC).

### 1.5 Surgery

Animals were prepared for head fixation by attaching two metal nuts to the skull using a construct made of a micro glass composite. The full procedure is described in van Alphen et al. (2009). Mice were given at least 3 days following surgery to recover before the start of any experimental paradigm.

### 1.6 Stimulus setup

Optokinetic stimuli were created using a modified Electrohome Marquee 9000 CRT projector (Christie Digital Systems, Cypress CA, USA) with a spatial resolution of at least 0.1 degrees and a temporal resolution of 0.01 s. The average luminance was kept constant at 17.5 cd/m^2^. The stimuli were projected via mirrors onto three transparent anthracite-colored screens (156*125 cm), which were placed in a triangular formation around the recording setup (Fig 2A). This created a green monochrome panoramic stimulus fully surrounding the animal. The stimuli were programmed in C++ and rendered in openGL. They each consisted of 1592 green dots (2 degrees diameter) equally spaced on a virtual sphere with its center at eye height above the center of the table. Moving stimuli were generated by rotating the virtual sphere around its vertical axis in sinusoidal patterns of different frequency and amplitude, so that all the dots moved coherently and in phase.

Vestibular stimulation was given by means of a motorized (Mavilor-DC motor 80, Mavilor Motors S.A., Barcelona, Spain) vestibular table that had its axis aligned with the center of the visual stimulus. The driving signal of both the visual and vestibular stimulation, which specified the required position, was computed and delivered by a CED Power1401 data acquisition interface (Cambridge Electronic Design, Cambridge, UK) with a resolution of 0.1 ° and 0.01 s.

### 1.7 Eye movement recordings

Mice were immobilized by placing them in a plastic tube, with the head pedestal bolted to a restrainer that allowed translations in three dimensions such that the eye of the mouse was placed in the center of the visual stimulus and thus above the rotation axis of the turn table, in front of the eye position recording camera.

Eye movements were recorded with an infrared video system (Iscan ETL-200, Iscan, Burlington, MA, USA). Images of the eye were captured at 120 Hz with an infrared sensitive CCD camera [see van Alphen et al. (2009) for more details]. To keep the field of view as free from obstacles as possible, the camera and lens were mounted under the table surface, and recordings were made with a hot mirror that was transparent to visible light and reflective to infrared light (Fig. 2B). The eye was illuminated with two infrared LEDs at the base of the hot mirror. The camera, mirror and LEDs were all mounted on an arm that could rotate about the vertical axis over a range of 26.1° (peak to peak). Eye movement recordings and calibration procedures were similar to those described by Stahl et al. (2000). Eye position was stored, along with the stimulus traces on hard disk for offline analysis.

### 1.8 Experimental Paradigms

#### 1.8.1 Optokinetic Reflex

The OKR (N=9) was tested using visual stimuli, while the mouse was kept stationary. We presented sinusoidal stimuli containing a wide range of frequencies (0.1, 0.2, 0.4, 0.8, 1.6 and 3.2 Hz) and amplitudes (0.5, 1.0, 2.0, 4.0, 6.0 and 8.0°), all about the earth vertical axis.

#### 1.8.2 Vestibulo-ocular Reflex

The VOR (N=9) was tested with vestibular stimulation in the dark. Stimulus amplitudes and frequencies were identical to those used for the OKR, except that stimuli with a peak velocity higher than 60 °/s were discarded, because of mechanical considerations. Again, only rotations about the vertical axis were made.

#### 1.8.3 Visually enhanced VOR and suppressed VOR

The vVOR (N=9) and the sVOR (N=6) protocols were identical to the VOR stimulation, except for the visual stimulation. During vVOR the visual stimulus was on, but stationary; during sVOR the visual stimulus was on and moved in phase and at the same amplitude as the turn table.

These four stimulus protocols were presented blockwise in 1 or 2 experimental sessions. Within each protocol the stimulus conditions were presented in random order to prevent effects of either learning or fatigue. All stimuli were presented for at least 5 cycles. The other protocols were performed separately.

#### 1.8.4 Non-periodic stimulation

For non-periodic stimulation we opted to give Sum-of-Sine (SoS) stimuli. In these SoS conditions, the two constituent frequencies were chosen that had no harmonic relation. Four SoS frequency combinations were used in this study: 0.6/0.8 Hz, 0.6/1.0 Hz, 0.8/1.0 Hz and 1.0/1.9 Hz. Amplitude was either one or two degrees for each frequency component. Either both frequencies had the same amplitude (both 1° or both 2°) or they had different amplitude (one at 1° and the other at 2°). This led to a total of 24 types of stimuli in each of the OKR, VOR, vVOR and sVOR SoS conditions. 8 mice were used in this paradigm and they all performed all conditions.

#### 1.8.5 Drift in the dark

In order to compute the plant time constant (see Supplementary Material, eq 15), we needed the mouse eye to drift in the dark from an eccentric position to the center of the oculomotor range. To do so, a visual scene moved slowly horizontal, thus making the eye move eccentrically. Subsequently, the light was turned off, and the mouse was in complete darkness. We then recorded the drift of the eye towards the center. By fitting an exponential function to this drift, the plant time constant was calculated. 6 mice were measured over a range of drift amplitudes between 4 and 10 degrees, the number of drift repetitions was on average around 6 per amplitude per mouse.

#### 1.8.6 VOR adaptation

VOR gain down adaptation (N=7) experiments consisted of 6 testing sessions and 5 training trials. Duration of each testing / training trial was 60s / 300s respectively. Sinusoidal (1 Hz, 5°) vestibular stimulation was applied in the dark for the testing sessions. During training sessions vestibular stimulation was accompanied by optokinetic sinusoidal stimulation of the same amplitude, phase and frequency (thus resulting in a stable head fixed visual surrounding).

### 1.9 Data Analysis

The Matlab (Matlab; The MathWorks, Natick, MA) code required for replication of the analysis presented in this paper is available on the Open Science Framework website (https://osf.io/feq7c/). Measured eye responses were analyzed offline. Position signals were transformed into velocity signals by a Savitski-Golay differentiating filter (cut-off frequency 50 Hz with a 3° polynomial) and were then smoothed with a median Gaussian filter (width 50 ms). Nystagmus fast phases and saccades were removed with a velocity threshold of 150°/s and with an FIR Butterworth low pass filter optimized to the stimulus frequency (cutoff at 3x stimulus frequency). There were two primary outcome measures in this study: gain and phase.

Gain and phase was extracted from the sinusoidal data by fitting a sinusoid and then using the gain and phase of the fit. The fit was done using a hierarchical Bayesian analysis using OpenBugs (Version 3.2.3, http://www.openbugs.net, [Lunn et al., 2009]). The precise details of the model used, as well as the parameters supplied to the OpenBugs algorithm, are provided below. In brief, the data for each trial for each mouse was assumed to be the result of a specific gain and phase specific to that trial, generated according to a distribution of gains and phases that were specific to the mouse. This distribution was, itself, generated according to hyper-parameters that characterize the population of mice. In addition, the noise in each trial was the result of a noise distribution characteristic of the mouse, which was generated according to hyper-parameters that characterized the population. Because our data was messy -- some mice had far more noise than others and some mice provided much more stable recording of eye movements than others -- the Bayesian approach allowed to incorporate all of the data in a robust manner, discounting the noisy or incomplete data when making estimates of the population parameters. Ultimately, we show the 95% high density intervals for the gain and phase of the individual mice in the bode plots (Figures 3-6B).

In order to summarize the mouse population in Figures 3-6A we generated 10,000 samples of posterior predictive mice. That is, for each of the 10,000 Bayesian samples, we selected an amplitude and phase according to the parameters for the mouse population, and then used that amplitude and phase to generate sinusoidal data. We used these 10,000 ‘typical’ mouse sinusoids to define a region of typical behavior. We characterized this region using the mean and standard deviation of these movements at each time step.

To summarize the similarity of the model response and the mouse population as a single value for each stimulus condition we employed Z-scores. Using the typical behavior we then calculated a Z-score by subtracting the model response at each time point from the center of the region of typical behavior and dividing by the standard deviation. This Z-score was then averaged across time points for each condition.

For the non-periodic data, gain and phase information were obtained by fitting two sine waves to the stimuli and the data in custom-made Matlab curve fitting routines using the least squares method.

For all experiments the fits of the sine waves to the eye movement data provided the amplitude and phase of the eye movements. The gain was calculated as the ratio of the amplitude of eye movement compared to the amplitude of the stimulus, phase was calculated by subtracting the phase of the stimulus from the movement. Thus, a positive phase value indicates a leading eye position signal.

## Statistics

Our statistics are geared to test whether the model behaves “similarly” to a typical mouse. This is different from the standard statistical test for effects and is also different from newly developed procedures that test for equivalence. We chose to test the confidence with which we could claim that model behavior lay within a “region of typical behavior” defined as the region within which 95% of mice are likely to fall. Thus, our p values represent the confidence with which we can make this statement.

For each condition, the gain and the phase of the model’s behavior were compared to the posterior predictive distribution of gains and phases of the mice. That is, for each Bayesian sample, we took the population mean and the population standard deviation for the gain. This gave us, for each Bayesian sample, an estimate of the mouse typical parameter value, from the mean minus 1.96 times the population standard deviation to the mean plus 1.96 times the population standard deviation. We determined the percentage of samples for which the value of the gain in the model lay within this typical region. We used this as a measure of the posterior predictive probability that our model gain was similar to those of a typical mouse. We used an identical procedure for the phase.

## Bayesian Fitting Procedure

The gains and phases of the single sine experimental data were estimated using a Bayesian fitting procedure using OpenBugs (version 3.2.3). The model used is specified in full form below:

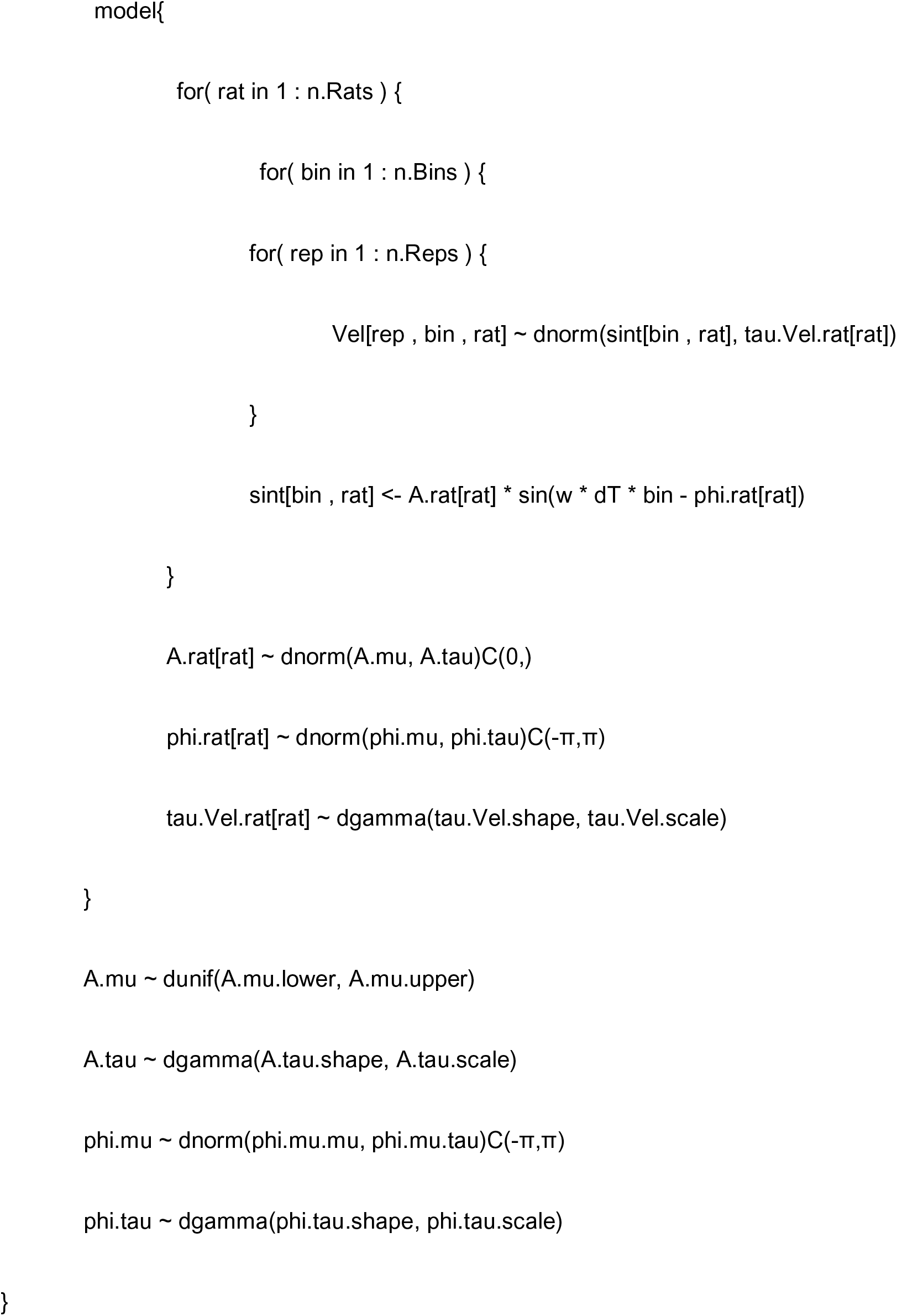

The fitting procedure was run with a burn-in of 500 samples, and then actual sampling of 10,000 samples in each of 3 chains. The initial values of the amplitude and phase of the fits were estimated from the data and each chain was initialized with a different precision (an order of magnitude between each). Convergence was assessed by manual inspection of the overlap of the chains and of the smoothness and overlap of the histograms for the posterior distribution of each parameter.

## Figure Supplements

**Figure 1-figure supplement 1.**
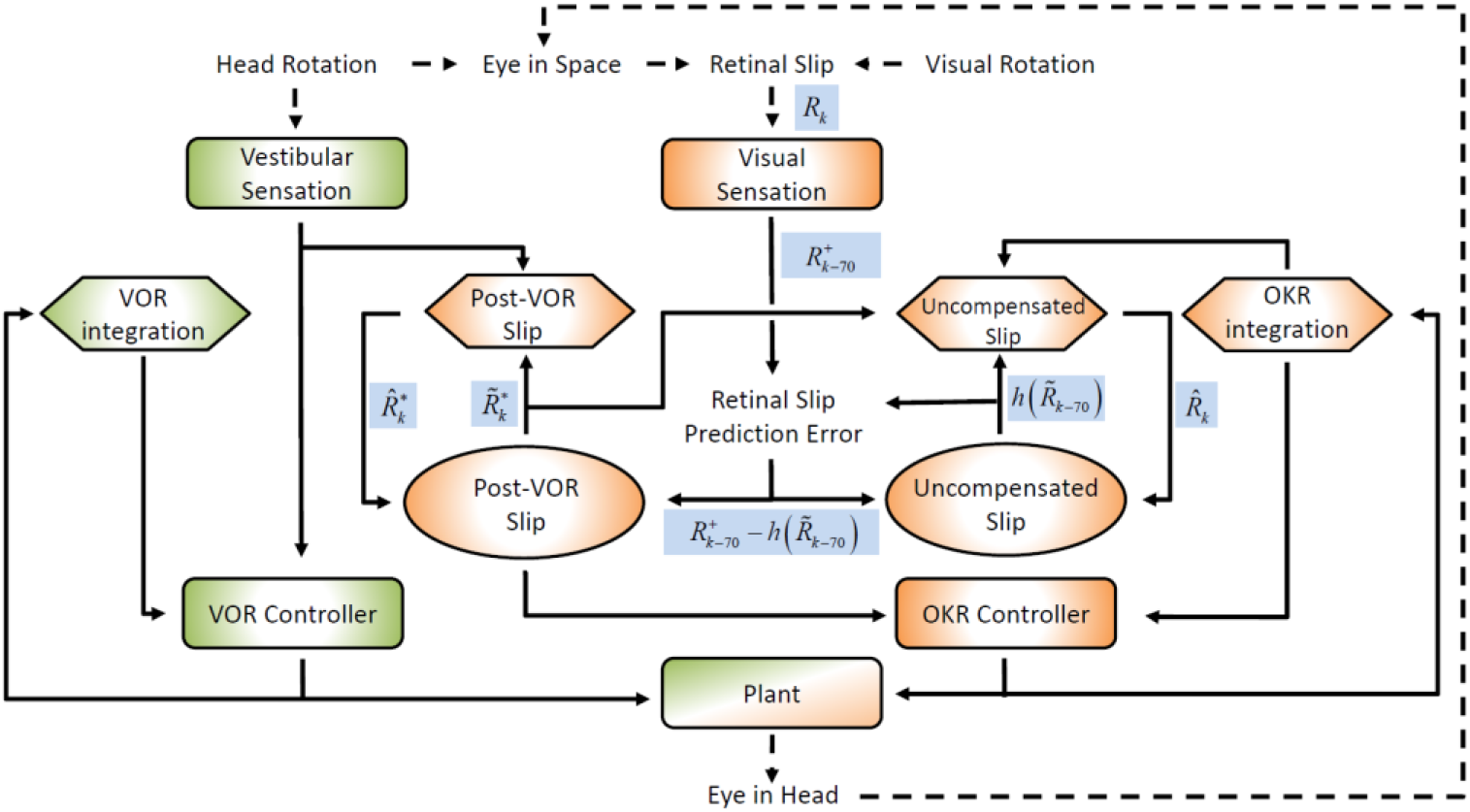
Schematic representation of the model architecture with the different internal and external representation of retinal slip indicated in blue rectangles adjacent to the corresponding arrows. The color and shape coding of the figure is maintained from Figure 1 in the main text.

**Figure 3-figure supplement 1.**
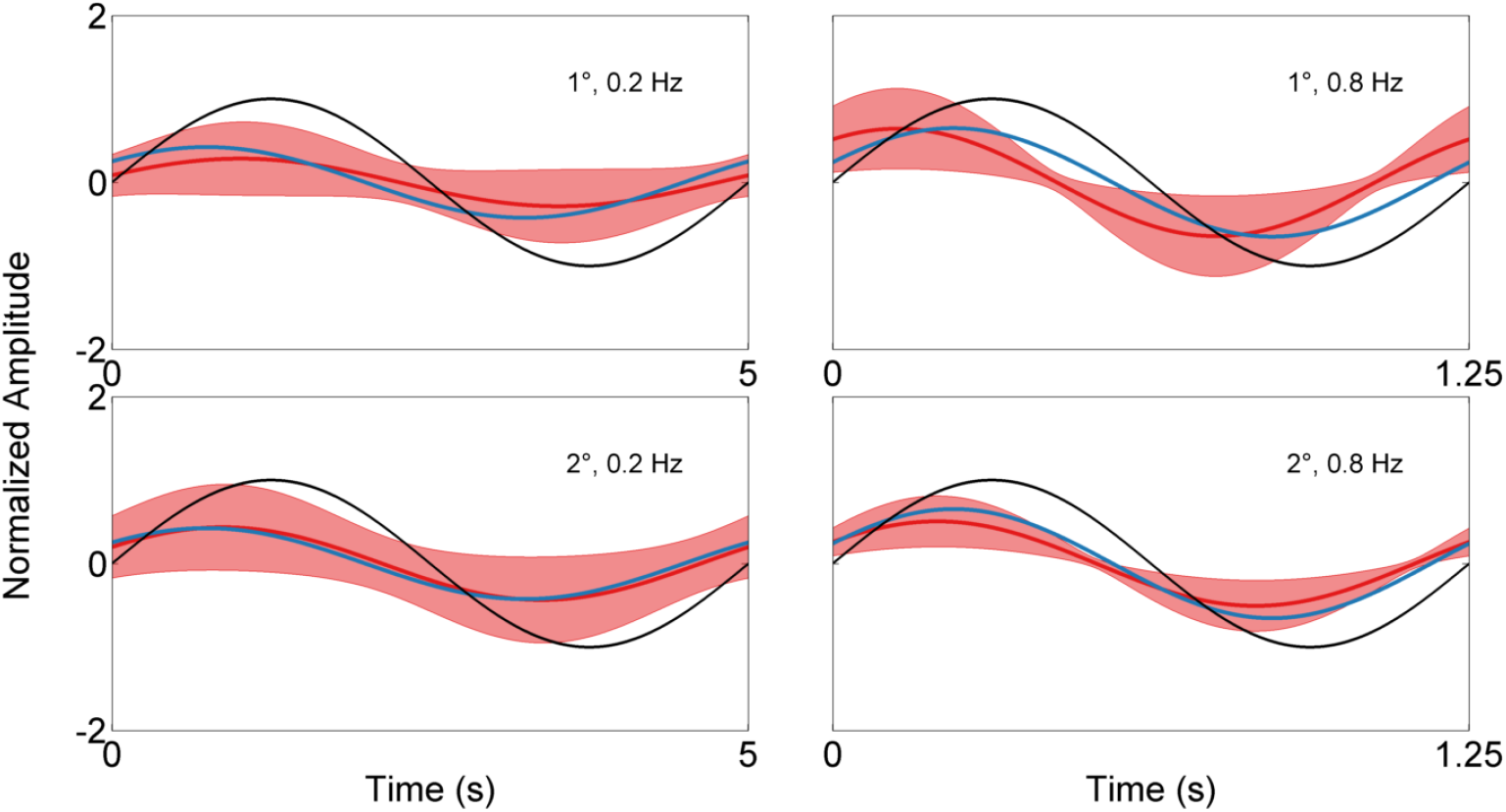
Examples of a sinusoid fitted to the model output (blue) and the mean measured response (red line) in response to a VOR stimulation (black line). The shaded red region represents the standard deviation of the population. The upper panels display the response to a 1° stimulus and the bottom panels correspond to a 2° stimulus. In both cases the left and right panels display the response to a 0.2 Hz and 0.8 Hz stimulus respectively. The format of the plots matches that of Figure 2 with the model output now represented with a fitted sinusoid (no longer including the high frequency noise included in the raw model output) for more direct comparison to the behavioral data which also represents the output of fitting sinusoids to the behavioral data.

**Figure 4-figure supplement 1.**
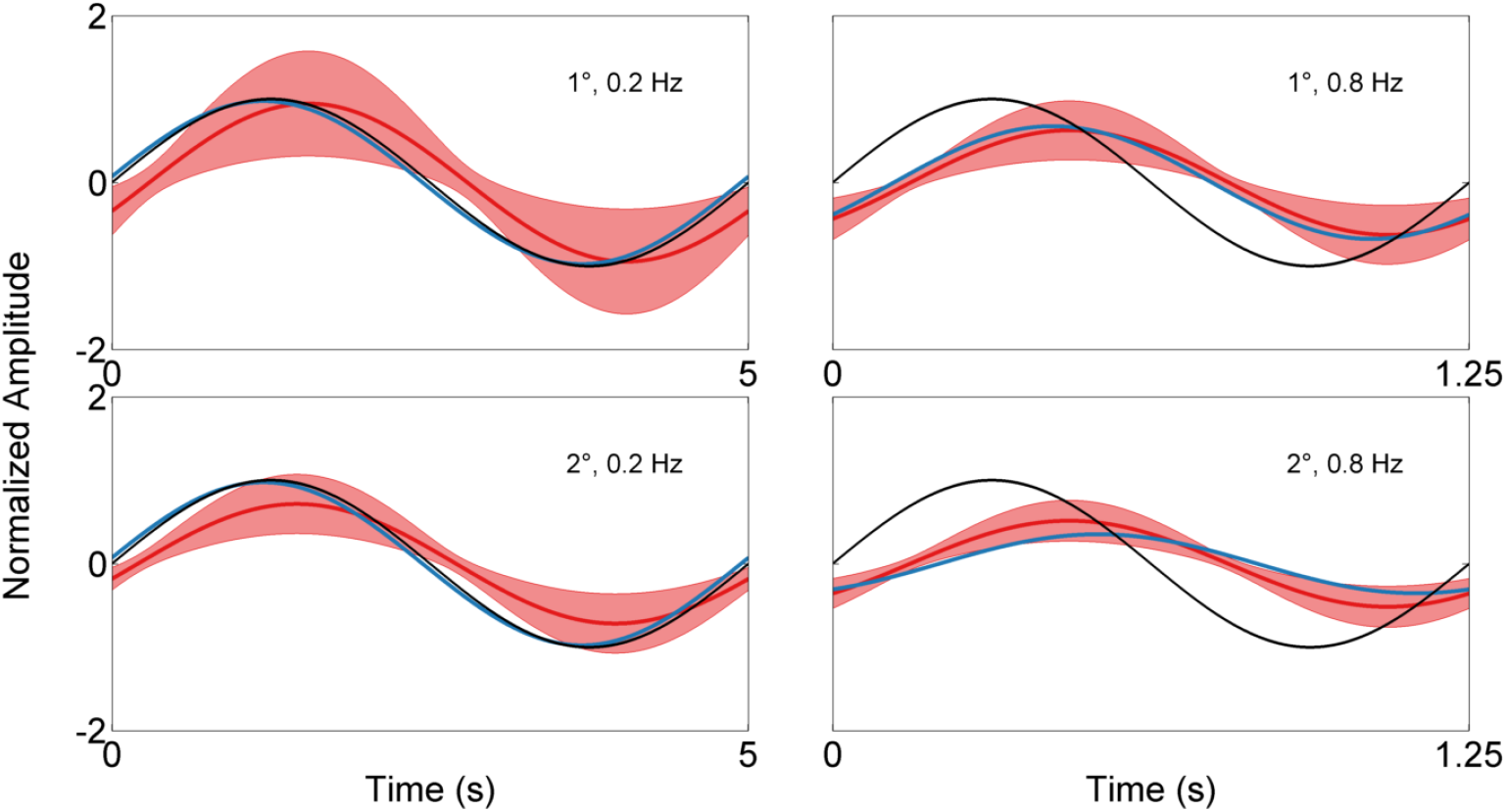
Examples of a sinusoid fitted to the model output (blue) and the mean measured response (red line) in response to a OKR stimulation (black line). The shaded red region represents the standard deviation of the population. The stimuli presented match that of Figure 4 and Figure 3-figure supplement 1.

**Figure 5-figure supplement 1.**
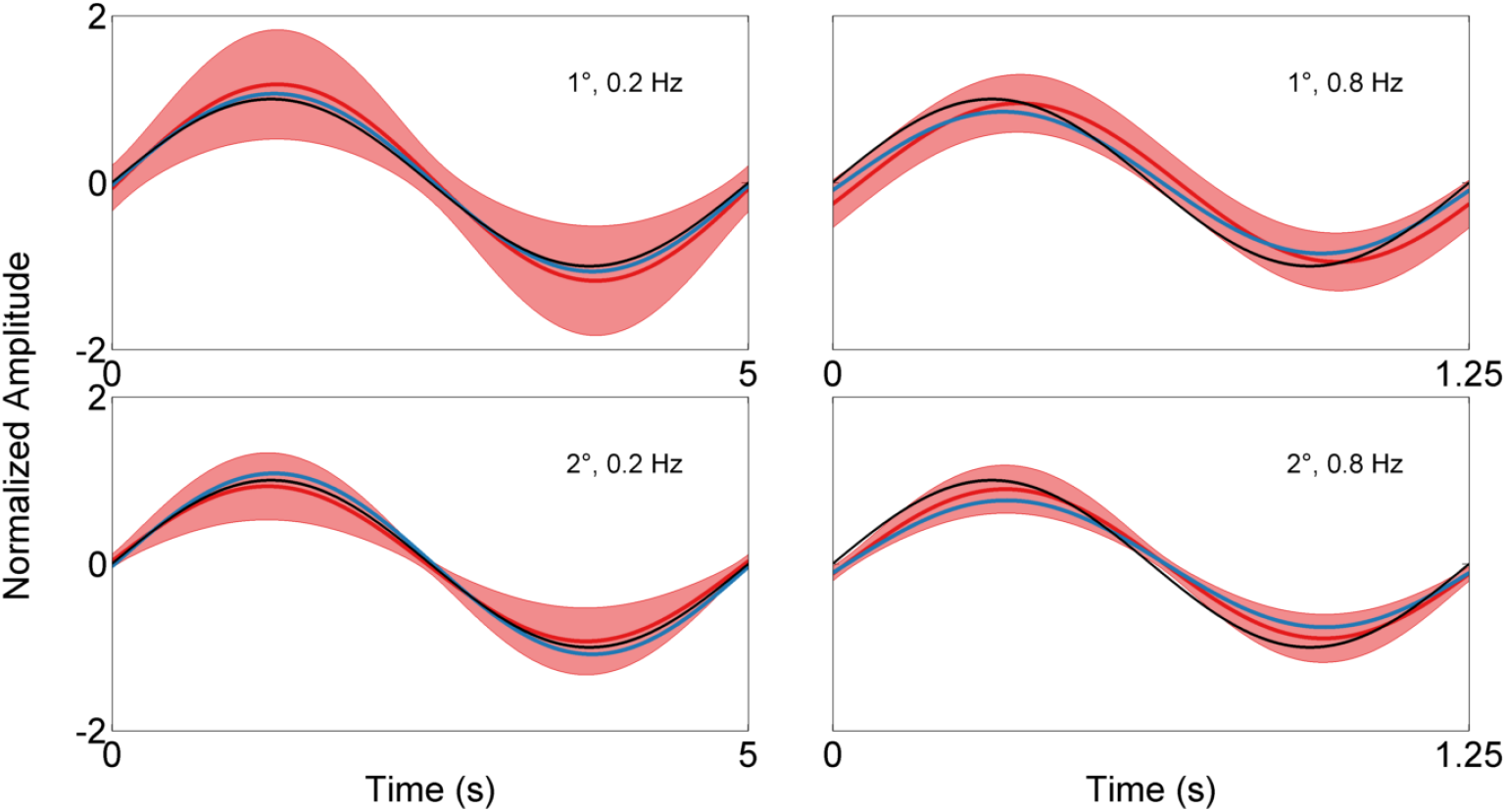
Examples of a sinusoid fitted to the model output (blue) and the mean measured response (red line) in response to a vVOR stimulation (black line). The shaded red region represents the standard deviation of the population. The stimuli presented match that of Figure 5 and Figure 3-figure supplement 1.

**Figure 6-figure supplement 1.**
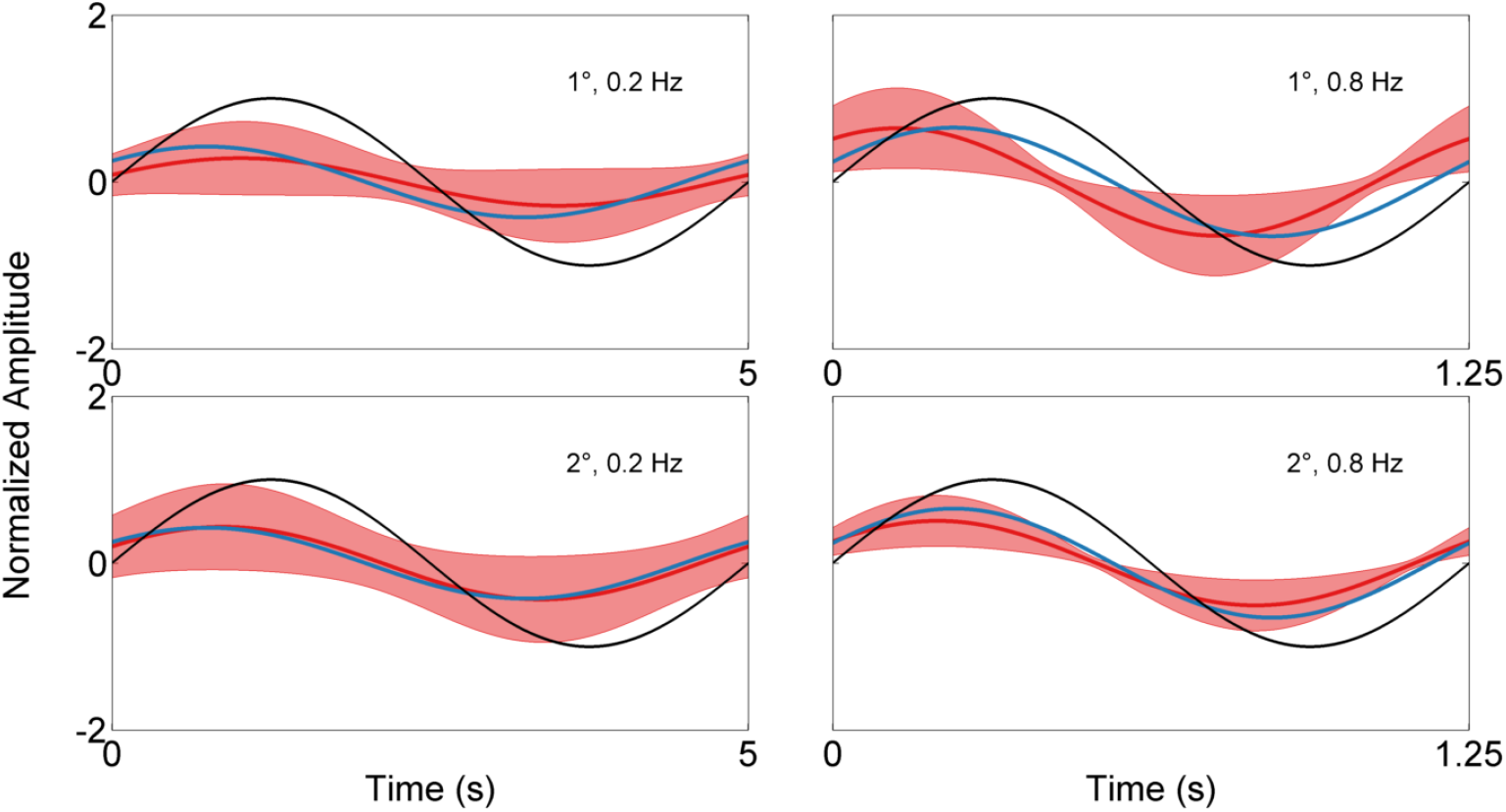
Examples of a sinusoid fitted to the model output (blue) and the mean measured response (red line) in response to a sVOR stimulation (black line). The shaded red region represents the standard deviation of the population. The stimuli presented match that of Figure 6 and Figure 3-figure supplement 1.

**Figure 7-figure supplement 1.**
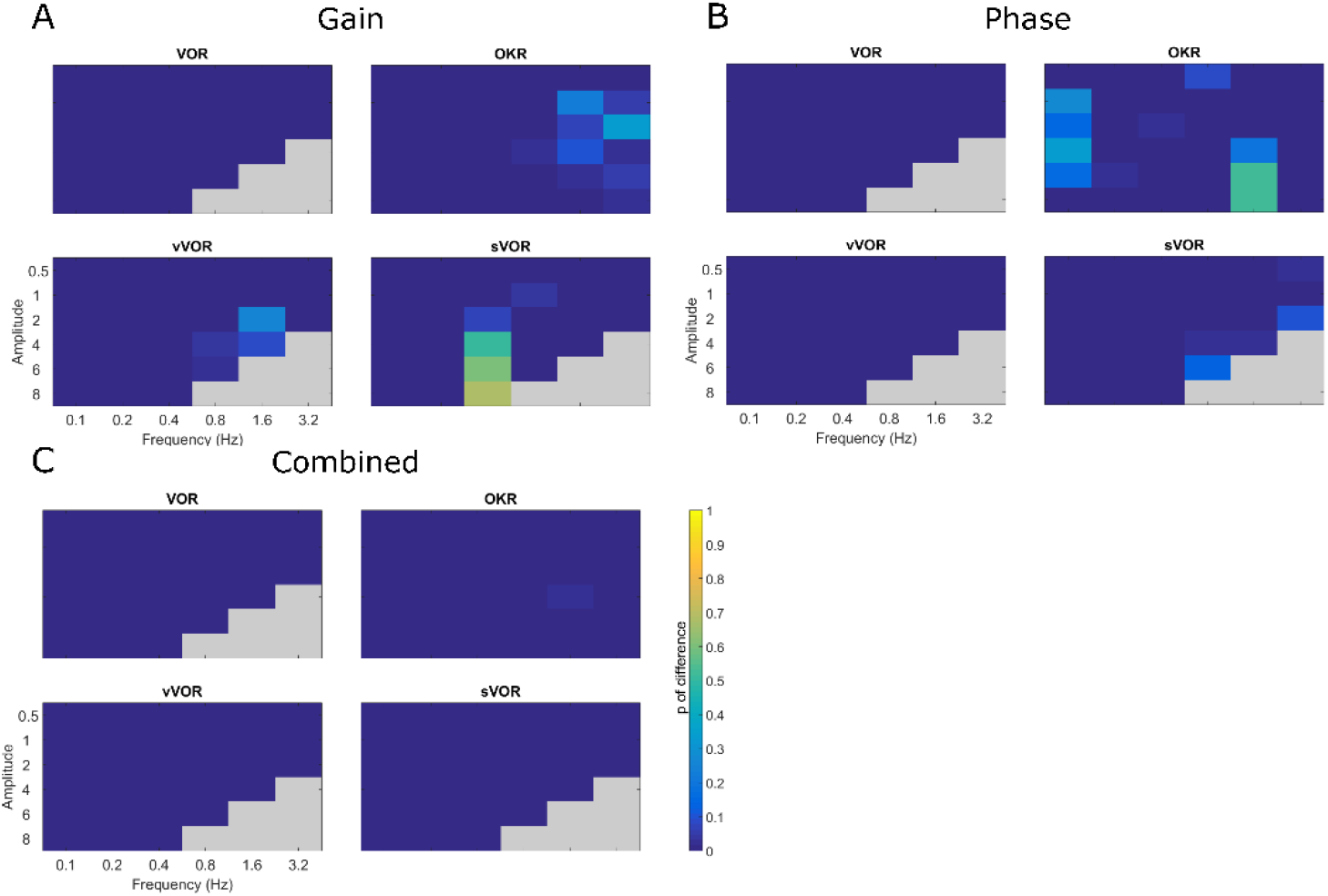
Summary of the statistical testing representing the probability that the model response falls outside the range of a ‘typical mouse’ in terms of gain (A), phase (B) and the combined probability (C). Overall, despite deviations in gain or phase in a minority of individual conditions, the model response is indistinguishable from the experimental data as indicated by the ‘cool’ colors in the combined probability graph. Full details of each individual test performed are available as tables in the source data for this figure.

**Figure 8-figure supplement 1.**
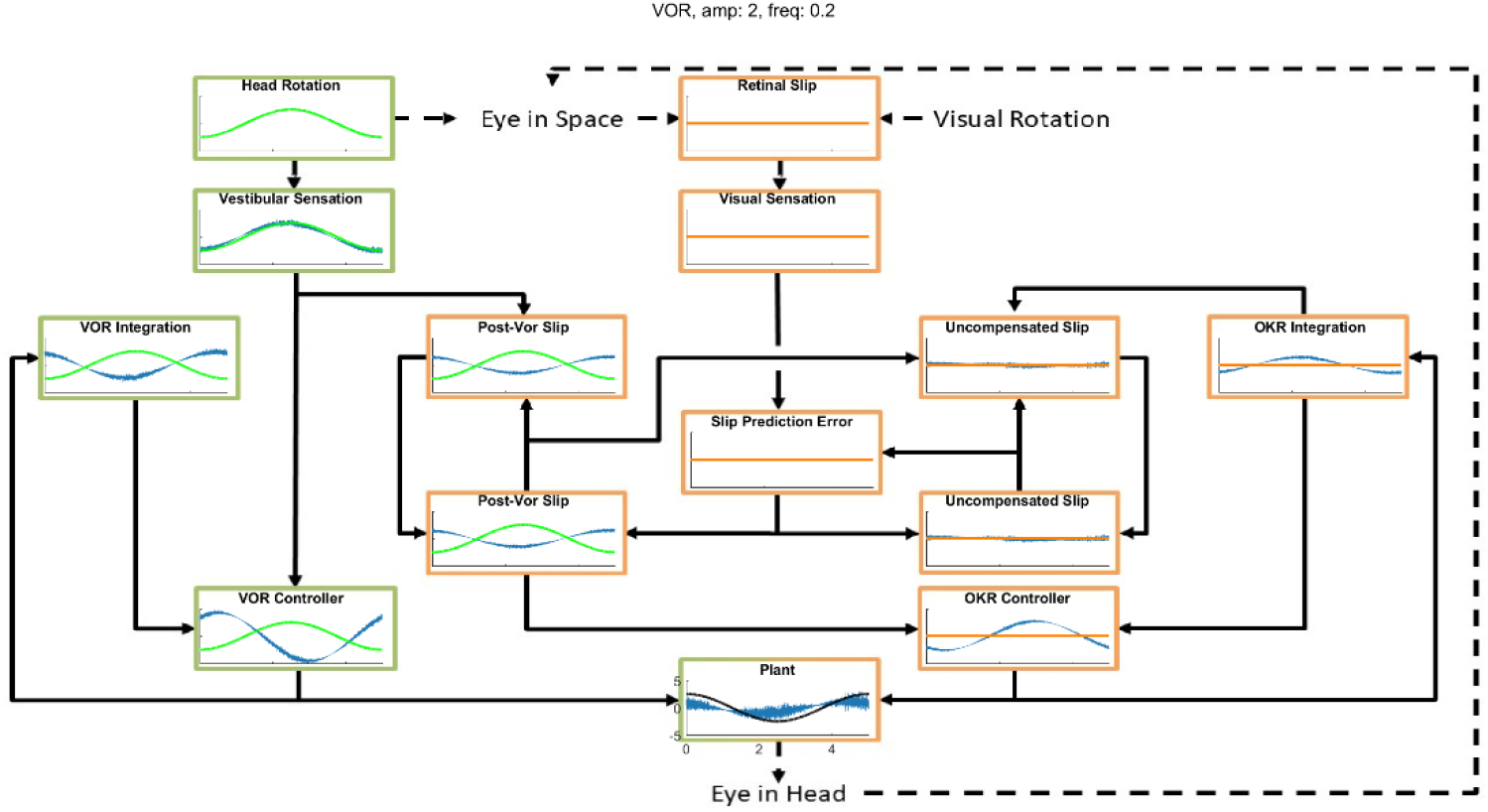
An example of the model dynamics for one cycle of the simulation in the VOR condition (Stimulation amplitude of 2 degrees at a frequency of 0.2 Hz) at a time by which the system has reached a steady state. The layout matches the model schematic presented in Figure 1. In each box the blue line represents the output of the computation performed, the green or orange line represents the appropriate stimulus, vestibular and visual respectively.

**Figure 8-figure supplement 2.**
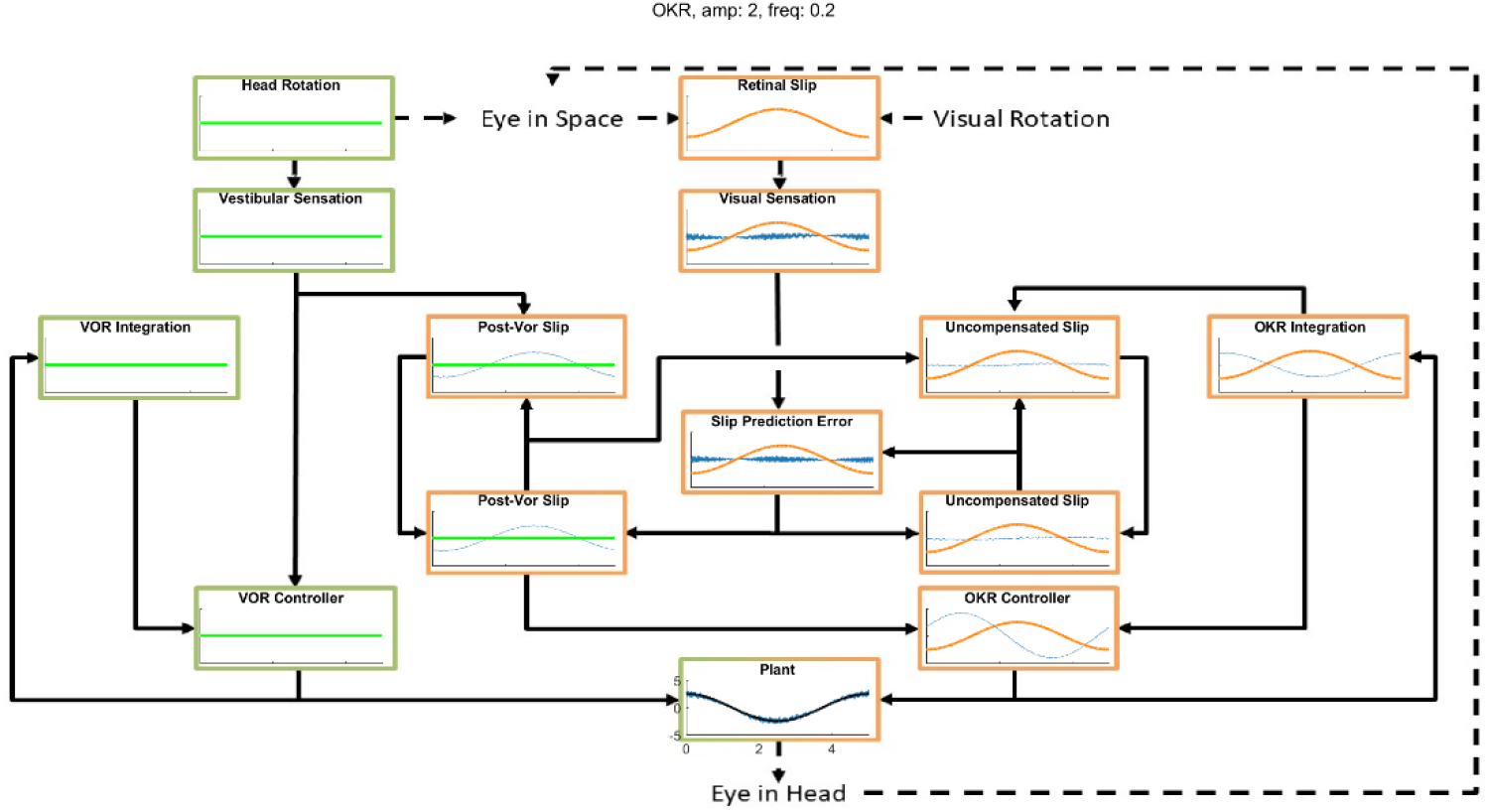
An example of the model dynamics for one cycle of the simulation in the OKR condition (Stimulation amplitude of 2 degrees at a frequency of 0.2 Hz) at a time by which the system has reached a steady state. The layout matches the model schematic presented in Figure 1. In each box the blue line represents the output of the computation performed, the green or orange line represents the appropriate stimulus, vestibular and visual respectively.

**Figure 9-figure supplement 1.**
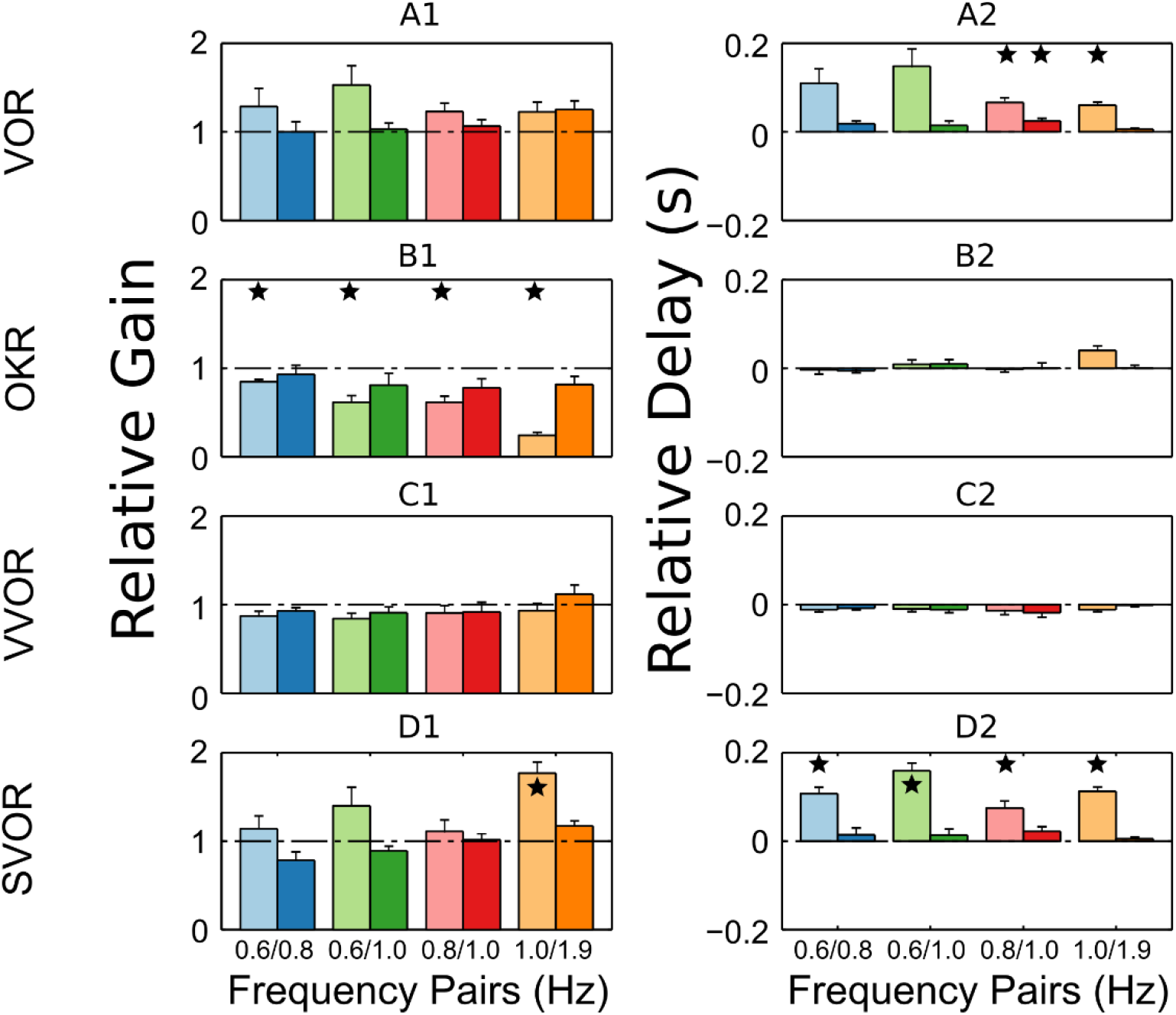
Summary of the behavioral response of c57BL/6 mince (n=8) to Sum of Sines stimulation. The response is described in terms of gains and lags relative to the gain and lag recorded in response to the single frequency component presented in isolation. A linear system will produce only relative gains of 1 and relative delays of 0, indicated by dashed horizontal lines on each plot. The pattern of nonlinearities produced is similar to that produced by the full model (Figure 7A). This figure is directly reproduced from Sibindi et al. (2016; Figure 6).

## References

van Alphen, A.M., Stahl, J.S., and De Zeeuw, C.I. (2001). The dynamic characteristics of the mouse horizontal vestibulo-ocular and optokinetic response. Brain Res. 890, 296–305.

van Alphen, A.M., Schepers, T., Luo, C., and De Zeeuw, C.I. (2002). Motor Performance and Motor Learning in Lurcher Mice. Ann. N. Y. Acad. Sci. 978, 413–424.

van Alphen, B., Winkelman, B.H.J., and Frens, M.A. (2009). Age- and Sex-Related Differences in Contrast Sensitivity in C57Bl/6 Mice. Investig. Opthalmology Vis. Sci. 50, 2451.

van Alphen, B., Winkelman, B.H.J., and Frens, M.A. (2010). Three-Dimensional Optokinetic Eye Movements in the C57BL/6J Mouse. Investig. Opthalmology Vis. Sci. 51, 623.

Belknap, D.B., and McCrea, R.A. (1988). Anatomical connections of the prepositus and abducens nuclei in the squirrel monkey. J. Comp. Neurol. 268, 13–28.

Belton, T., and McCrea, R.A. (2000). Role of the Cerebellar Flocculus Region in Cancellation of the VOR During Passive Whole Body Rotation. J. Neurophysiol. 84, 1599–1613.

Blazquez, P., Hirata, Y., and Highstein, S. (2004). The vestibulo-ocular reflex as a model system for motor learning: what is the role of the cerebellum? The Cerebellum 3, 188–192.

Büttner-Ennever, J.A., and Büttner, U. (1992). Neuroanatomy of the ocular motor pathways. Baillières Clin. Neurol. 1, 263–287.

Cannon, S.C., and Robinson, D.A. (1987). Loss of the neural integrator of the oculomotor system from brain stem lesions in monkey. J. Neurophysiol. 57, 1383–1409.

Cheron, G., Gillis, P., and Godaux, E. (1986a). Lesions in the cat prepositus complex: effects on the optokinetic system. J. Physiol. 372, 95–111.

Cheron, G., Godaux, E., Laune, J.M., and Vanderkelen, B. (1986b). Lesions in the cat prepositus complex: effects on the vestibulo-ocular reflex and saccades. J. Physiol. 372, 75–94.

Clopath, C., Badura, A., Zeeuw, C.I.D., and Brunel, N. (2014). A Cerebellar Learning Model of Vestibulo-Ocular Reflex Adaptation in Wild-Type and Mutant Mice. J. Neurosci. 34, 7203–7215.

Collewijn, H. (1969). Optokinetic eye movements in the rabbit: Input-output relations. Vision Res. 9, 117–132.

Delgado-García, J.M. (2000). Why move the eyes if we can move the head? Brain Res. Bull. 52, 475–482.

Faulstich, B.M., Onori, K.A., and du Lac, S. (2004). Comparison of plasticity and development of mouse optokinetic and vestibulo-ocular reflexes suggests differential gain control mechanisms. Vision Res. 44, 3419–3427.

Frens, M.A., and Donchin, O. (2009). Forward Models and State Estimation in Compensatory Eye Movements. Front. Cell. Neurosci. 3.

Gao, Z., van Beugen, B.J., and De Zeeuw, C.I. (2012). Distributed synergistic plasticity and cerebellar learning. Nat. Rev. Neurosci. 13, 619–635.

Gerrits, N.M., Epema, A.H., and Voogd, J. (1984). The mossy fiber projection of the nucleus reticularis tegmenti pontis to the flocculus and adjacent ventral paraflocculus in the cat. Neuroscience 11, 627–644.

Ghasia, F.F., Meng, H., and Angelaki, D.E. (2008). Neural Correlates of Forward and Inverse Models for Eye Movements: Evidence from Three-Dimensional Kinematics. J. Neurosci. 28, 5082– 5087.

Glasauer, S. (2007). Current models of the ocular motor system. Dev. Ophthalmol. 40, 158–174.

Glickstein, M., Gerrits, N., Kralj-Hans, I., Mercier, B., Stein, J., and Voogd, J. (1994). Visual pontocerebellar projections in the macaque. J. Comp. Neurol. 349, 51–72.

Graf, W., Simpson, J.I., and Leonard, C.S. (1988). Spatial organization of visual messages of the rabbit’s cerebellar flocculus. II. Complex and simple spike responses of Purkinje cells. J. Neurophysiol. 60, 2091–2121.

Green, A.M., Meng, H., and Angelaki, D.E. (2007). A Reevaluation of the Inverse Dynamic Model for Eye Movements. J. Neurosci. 27, 1346–1355.

Haar, S., and Donchin, O. A revised computational neuroanatomy for motor control.

Haith, A., and Vijayakumar, S. (2007). Robustness of VOR and OKR adaptation under kinematics and dynamics transformations. In 2007 IEEE 6th International Conference on Development and Learning, pp. 37–42.

Jordan, M.I., and Rumelhart, D.E. (1992). Forward Models: Supervised Learning with a Distal Teacher. Cogn. Sci. 16, 307–354.

Kawato, M., and Gomi, H. (1992). The cerebellum and VOR/OKR learning models. Trends Neurosci. 15, 445–453.

Langer, T., Fuchs, A.F., Scudder, C.A., and Chubb, M.C. (1985). Afferents to the flocculus of the cerebellum in the rhesus macaque as revealed by retrograde transport of horseradish peroxidase. J. Comp. Neurol. 235, 1–25.

Lisberger, S.G. (2009). Internal models of eye movement in the floccular complex of the monkey cerebellum. Neuroscience 162, 763–776.

Lunn, D., Spiegelhalter, D., Thomas, A., and Best, N. (2009). The BUGS project: Evolution, critique and future directions. Stat. Med. 28, 3049–3067.

McCrea, R.A., and Baker, R. (1985). Anatomical connections of the nucleus prepositus of the cat. J. Comp. Neurol. 237, 377–407.

Oyster, C.W., Takahashi, E., and Collewijn, H. (1972). Direction-selective retinal ganglion cells and control of optokinetic nystagmus in the rabbit. Vision Res. 12, 183–193.

Parrell, B., Ramanarayanan, V., Nagarajan, S., and Houde, J. (2019). The FACTS model of speech motor control: fusing state estimation and task-based control. BioRxiv 543728.

Porrill, J., and Dean, P. (2007). Cerebellar Motor Learning: When Is Cortical Plasticity Not Enough? PLoS Comput Biol 3, e197.

Porrill, J., Dean, P., and Stone, J.V. (2004). Recurrent cerebellar architecture solves the motor-error problem. Proc. R. Soc. B Biol. Sci. 271, 789–796.

Rambold, H., Churchland, A., Selig, Y., Jasmin, L., and Lisberger, S.G. (2002). Partial Ablations of the Flocculus and Ventral Paraflocculus in Monkeys Cause Linked Deficits in Smooth Pursuit Eye Movements and Adaptive Modification of the VOR. J. Neurophysiol. 87, 912–924.

Robinson, D.A. (1981). The Use of Control Systems Analysis in the Neurophysiology of Eye Movements. Annu. Rev. Neurosci. 4, 463–503.

Sağlam, M., Lehnen, N., and Glasauer, S. (2011). Optimal Control of Natural Eye-Head Movements Minimizes the Impact of Noise. J. Neurosci. 31, 16185–16193.

Sağlam, M., Glasauer, S., and Lehnen, N. (2014). Vestibular and cerebellar contribution to gaze optimality. Brain 137, 1080–1094.

Schonewille, M., Belmeguenai, A., Koekkoek, S.K., Houtman, S.H., Boele, H.J., van Beugen, B.J., Gao, Z., Badura, A., Ohtsuki, G., Amerika, W.E., et al. (2010). Purkinje Cell-Specific Knockout of the Protein Phosphatase PP2B Impairs Potentiation and Cerebellar Motor Learning. Neuron 67, 618– 628.

Schonewille, M., Gao, Z., Boele, H.-J., Vinueza Veloz, M.F., Amerika, W.E., Šimek, A.A.M., De Jeu, M.T., Steinberg, J.P., Takamiya, K., Hoebeek, F.E., et al. (2011). Reevaluating the Role of LTD in Cerebellar Motor Learning. Neuron 70, 43–50.

Shadmehr, R., and Krakauer, J.W. (2008). A computational neuroanatomy for motor control. Exp. Brain Res. 185, 359–381.

Shin, S.-L., Zhao, G.Q., and Raymond, J.L. (2014). Signals and Learning Rules Guiding Oculomotor Plasticity. J. Neurosci. 34, 10635–10644.

Sibindi, T.M., Holland, P.J., van der Geest, J.N., Donchin, O., and Frens, M.A. (2016). Superposition Violations in the Compensatory Eye Movement System. Investig. Opthalmology Vis. Sci. 57, 3554.

Sohmer, H., Elidan, J., Plotnik, M., Freeman, S., Sockalingam, R., Berkowitz, Z., and Mager, M. (1999). Effect of noise on the vestibular system - Vestibular evoked potential studies in rats. Noise Health 2, 41.

Soodak, R.E., and Simpson, J.I. (1988). The accessory optic system of rabbit. I. Basic visual response properties. J. Neurophysiol. 60, 2037–2054.

Stahl, J.S., and Simpson, J.I. (1995). Dynamics of abducens nucleus neurons in the awake rabbit. J Neurophysiol 73, 1383–1395.

Stahl, J.S., van Alphen, A.M., and De Zeeuw, C.I. (2000). A comparison of video and magnetic search coil recordings of mouse eye movements. J. Neurosci. Methods 99, 101–110.

Stahl, J.S., James, R.A., Oommen, B.S., Hoebeek, F.E., and Zeeuw, C.I.D. (2006). Eye Movements of the Murine P/Q Calcium Channel Mutant Tottering, and the Impact of Aging. J. Neurophysiol. 95, 1588–1607.

Stahl, J.S., Thumser, Z.C., May, P.J., Andrade, F.H., Anderson, S.R., and Dean, P. (2015). Mechanics of mouse ocular motor plant quantified by optogenetic techniques. J. Neurophysiol. 114, 1455–1467.

Takemori, S., and Cohen, B. (1974). Loss of visual suppression of vestibular nystagmus after flocculus lesions. Brain Res. 72, 213–224.

Todorov, E., and Jordan, M.I. (2002). Optimal feedback control as a theory of motor coordination. Nat. Neurosci. 5, 1226–1235.

Wakita, R., Tanabe, S., Tabei, K., Funaki, A., Inoshita, T., and Hirano, T. (2017). Differential regulations of vestibulo-ocular reflex and optokinetic response by β- and α2-adrenergic receptors in the cerebellar flocculus. Sci. Rep. 7, 3944.

Winkelman, B., and Frens, M. (2006). Motor Coding in Floccular Climbing Fibers. J. Neurophysiol. 95, 2342–2351.

Yang, A., and Hullar, T.E. (2007). Relationship of Semicircular Canal Size to Vestibular-Nerve Afferent Sensitivity in Mammals. J. Neurophysiol. 98, 3197–3205.

Zee, D.S., Yamazaki, A., Butler, P.H., and Gucer, G. (1981). Effects of ablation of flocculus and paraflocculus of eye movements in primate. J. Neurophysiol. 46, 878–899.

## References

Åström, K.J., and Murray, R.M. (2008). Feedback Systems: An Introduction for Scientists and Engineers (Princeton, NJ: Princeton University Press).

Bradtke, S.J. (1993). Reinforcement Learning Applied to Linear Quadratic Regulation. In In Advances in Neural Information Processing Systems 5, (Morgan Kaufmann), pp. 295–302.

Burns, J.A., and Ou, Y.-R. (1994). Feedback control of the driven cavity problem using LQR designs. In, Proceedings of the 33rd IEEE Conference on Decision and Control, 1994, pp. 289–294 vol.1.

Frens, M.A., and Donchin, O. (2009). Forward models and state estimation in compensatory eye movements. Front. Cell. Neurosci. 3, 13.

Harris, C.M., and Wolpert, D.M. (1998). Signal-dependent noise determines motor planning. Nature 394, 780–784.

Lancaster, P., and Rodman, L. (1995). Algebraic Riccati Equations (Clarendon Press).

Lopez-Martinez, M., Diaz, J.M., Ortega, M.G., and Rubio, F.R. (2004). Control of a laboratory helicopter using switched 2-step feedback linearization. In American Control Conference, 2004. Proceedings of the 2004, pp. 4330–4335 vol.5.

Porrill, J., Dean, P., and Anderson, S.R. (2013). Adaptive filters and internal models: Multilevel description of cerebellar function. Neural Netw. 47, 134–149.

Stahl, J.S., and Simpson, J.I. (1995). Dynamics of abducens nucleus neurons in the awake rabbit. Neurophysiol. 73, 1383–1395.

Stahl, J.S., Thumser, Z.C., May, P.J., Andrade, F.H., Anderson, S.R., and Dean, P. (2015). Mechanics of mouse ocular motor plant quantified by optogenetic techniques. J. Neurophysiol. 114, 1455– 1467.

Todorov, E. (2004). Optimality principles in sensorimotor control (review). Nat. Neurosci. 7, 907–915.

Yang, A., and Hullar, T.E. (2007). Relationship of semicircular canal size to vestibular-nerve afferent sensitivity in mammals. J. Neurophysiol. 98, 3197–3205.

Yoshida, K., Watanabe, D., Ishikane, H., Tachibana, M., Pastan, I., and Nakanishi, S. (2001). A Key Role of Starburst Amacrine Cells in Originating Retinal Directional Selectivity and Optokinetic Eye Movement. Neuron 30, 771–780.

